# *In silico* modeling and simulation of neuroendocrine-immune modulation through adrenergic and 17β-estradiol receptors in lymphocytes show differential activation of cyclic adenosine monophosphate (cAMP)

**DOI:** 10.1101/353185

**Authors:** H. P. Priyanka, A. Thiyagaraj, R.S. Nair, G. Krithika, L. Hima, W. Hopper, S. ThyagaRajan

## Abstract

Sympathetic innervation of lymphoid organs and presence of 17β-estradiol (estrogen or E_2_) and adrenergic receptors (ARs) on lymphocytes suggests that sympathetic stimulation and hormonal activation may influence immune functions. Simulation of these pathways may help to understand the dynamics of neuroendocrine-immune modulation at the cellular and molecular level.

Dose- and receptor-dependent effects of 17β-estradiol and AR sub-type-specific agonists were established in vitro on lymphocytes from young male Sprague-Dawley rats and modeled in silico using MATLAB Simbiology toolbox. Kinetic principles were assigned to define receptor-ligand dynamics and concentration/time plots were obtained using Ode15s solvers at different time intervals for key regulatory molecules. Comparisons were drawn between in silico and *in vitro* data for validating the constructed model with sensitivity analysis of key regulatory molecules to assess their individual impacts on the dynamics of the system.

Adrenergic activation triggered pro-apoptotic signals while 17β-estradiol enhanced survival signals showing contradictory effects as observed in vitro. Treatment of lymphocytes with 17β-estradiol shows ten-fold increase in survival signals in a dose-dependent manner. cAMP (cyclic adenosine monophosphate) activation is crucial for the activation of survival signals through p-ERK (Extracellular Signal-Regulated Kinase) and p-CREB (cAMP Responsive Element Binding) protein.

Thus, the cross-talk between 17β-estradiol and adrenergic signaling pathways determines lymphocyte functions in a receptor subtype- and co-activation-dependent manner in health and disease.

## 1.0 Introduction

The neuroendocrine-immune network is a complex inter-regulatory system with wide plasticity in order to maintain systemic homeostasis [1-3]. In females, the cyclic fluctuations in the levels of gonadal hormones especially, 17β-estradiol affect the functioning of immune effector cells by binding to specific estrogen receptors [ER; 4-7].

In the periphery, *in vitro* and *in vivo* 17β-estradiol-stimulation has been shown to enhance splenocyte proliferation and cytokine production through the alteration of specific signaling molecules [2, 4, 8]. Estrogen enhances splenocyte proliferation, Interferon-γ (IFN-γ) expression, through p-ERK and p-CREB signals, enhances activity of antioxidant enzymes (AOE) including superoxide dismutase (SOD) and catalase (CAT) and increases splenocyte nitric oxide (NO) expression dose dependently [4]. In the resting state, the close apposition of T-lymphocytes in direct synaptic association with sympathetic noradrenergic (NA) nerve fibers that innervate the lymphoid organs renders them highly responsive to norepinephrine [NE; 7, 9-13]. Since both 17β-estradiol and NE mediate their effects on lymphocytes through their specific receptors, their down-stream effects are dependent upon the kinetic parameters that govern receptor-ligand interactions [10, 14-16]. Previous studies from our laboratory and others have shown that both 17β-estradiol and norepinephrine signaling cascades involve similar molecules (p-ERK, p-CREB, cAMP and p-Akt) in modulating the expression of similar cytokines (IFN-γ and interleukin-2 (IL-2)) thereby, altering cell-mediated immune functions [2, 4, 17-18]. Thus, the scope of cross-talk between the two pathways during multiple ligand-receptor interactions is complex. Considering that NE can bind to α1-/α2- or β1-/β2-/β3-adrenoceptors (AR) on lymphocytes and 17β-estradiol can bind to cytosolic or nuclear 17β-estradiol receptors (cER and nER) and initiate signals that target similar down-stream signaling molecules, the underlying kinetic parameters can influence the outcome of the cross-talk through synergistic, additive or diminutive effects.

Our lab has delineated these effects in splenocytes using adrenergic receptor-specific agonists and antagonists in the presence and absence of estrogen [17, 18]. α1-AR agonist Phenylephrine enhances p-ERK and p-CREB expression, while α2-AR agonist clonidine enhances p-Akt expression and may play a crucial role in the sustenance of naïve cells in the secondary lymphoid organs [17]. β2-AR agonist terbutaline enhances the expression of p-ERK, p-CREB and p-Akt signals, antioxidant enzyme activities including CAT and SOD and the production of NO in splenocytes [18]. In this study, we have superimposed the four signaling cascades (Estrogen signaling, α1-AR signaling, α2-AR signaling and β2-AR signaling) from our in vitro studies, and created a dynamic model of resting lymphocyte functions. In order to simulate real time events we have used kinetic principles that govern receptor-ligand activation based on secondary data from available literature along with concentration data from *in vitro* studies published from our laboratory [4,17,18]. The present study aims to create a computational model of the pathways studied in vitro so that the scope of the intercommunication between these pathways can be better understood in health and disease.

## 2.0 Methods

### 2.1 Model building

The model was built using both primary and secondary data sources. The dose- and receptor-dependent effects of 17β-estradiol and AR sub-type-specific agonists were established *in vitro* on lymphocytes from young male Sprague-Dawley rats in previous studies published by our laboratory [17,18]. Briefly, splenic lymphocytes were isolated from young male Sprague-Dawley rats and treated with different doses of Estrogen [4], AR-α agonists phenylephrine and clonidine [17] and AR-β agonist terbutaline [18] for a period of 24-72 hours. In these studies, the concentration of the signaling molecules, cytokines, and other secondary molecules was determined using ELISA and these data were used to construct the model (Tables 1, 2). The possible cross-talk pathway was mapped using receptor-specific inhibitors and inhibitors of signal transduction molecules and was modeled *in silico* using the MATLAB Simbiology toolbox [MATLAB and SimBiology toolbox Release 2019a, The Mathworks, inc., Natick, Massachusetts, United States]. Receptor-ligand dynamics were defined based on kinetic principles from available literature and previous studies from our laboratory [4, 17,18]. State versus time plots were obtained using Ode15s solvers at 24 hours or 72 hrs for long-term effects on lymphocyte functions. Comparisons were drawn between *in silico* and *in vitro* data for validating the constructed model. Sensitivity analysis was performed on key regulatory molecules to assess their individual impacts on the dynamics of the system. The model was uploaded in Github database.

(https://github.com/Anandt1082/Neuroimmunomodulation-by-estrogen-and-adrenergic-agonists.git).

**Table 1:**
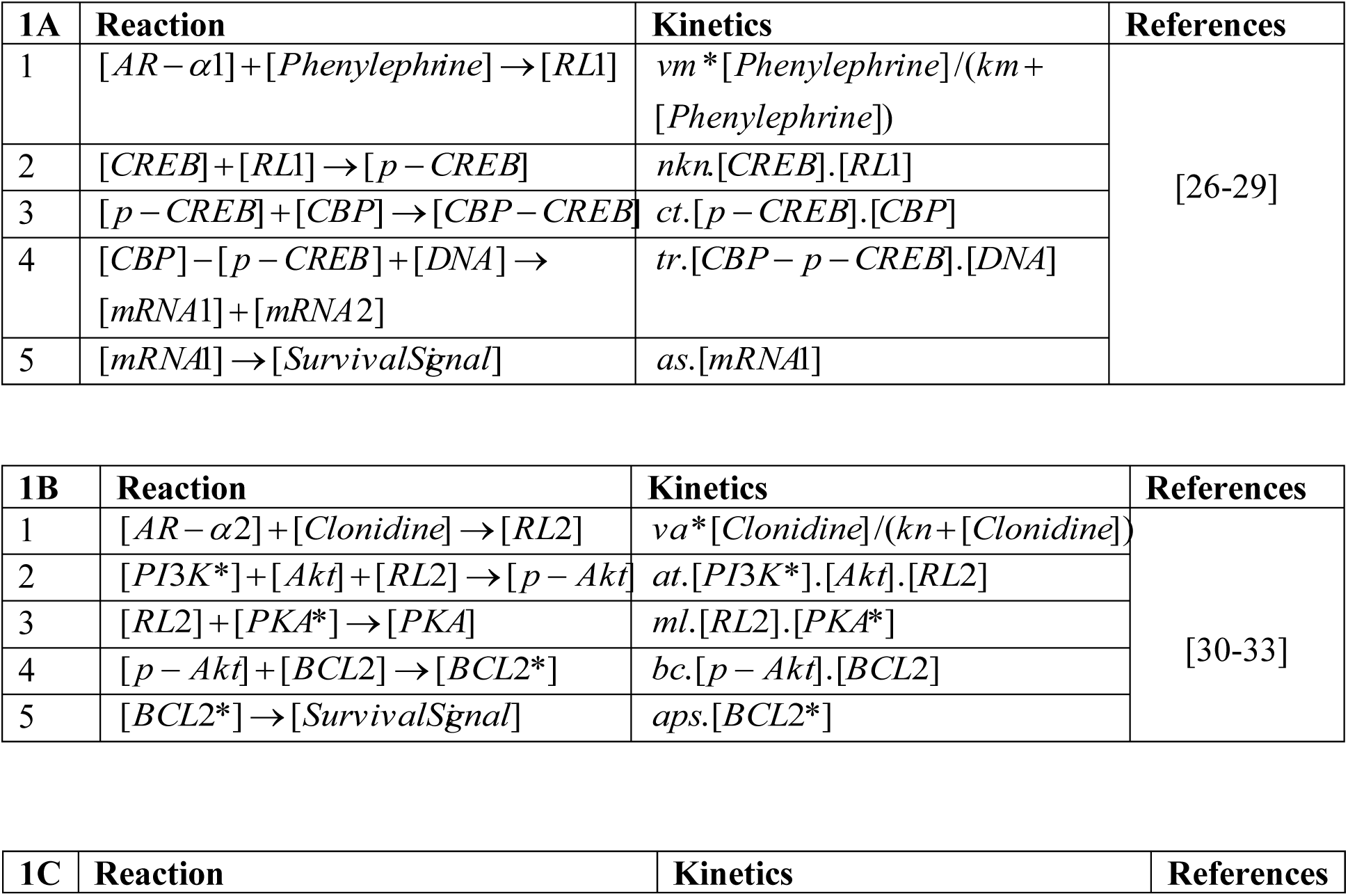
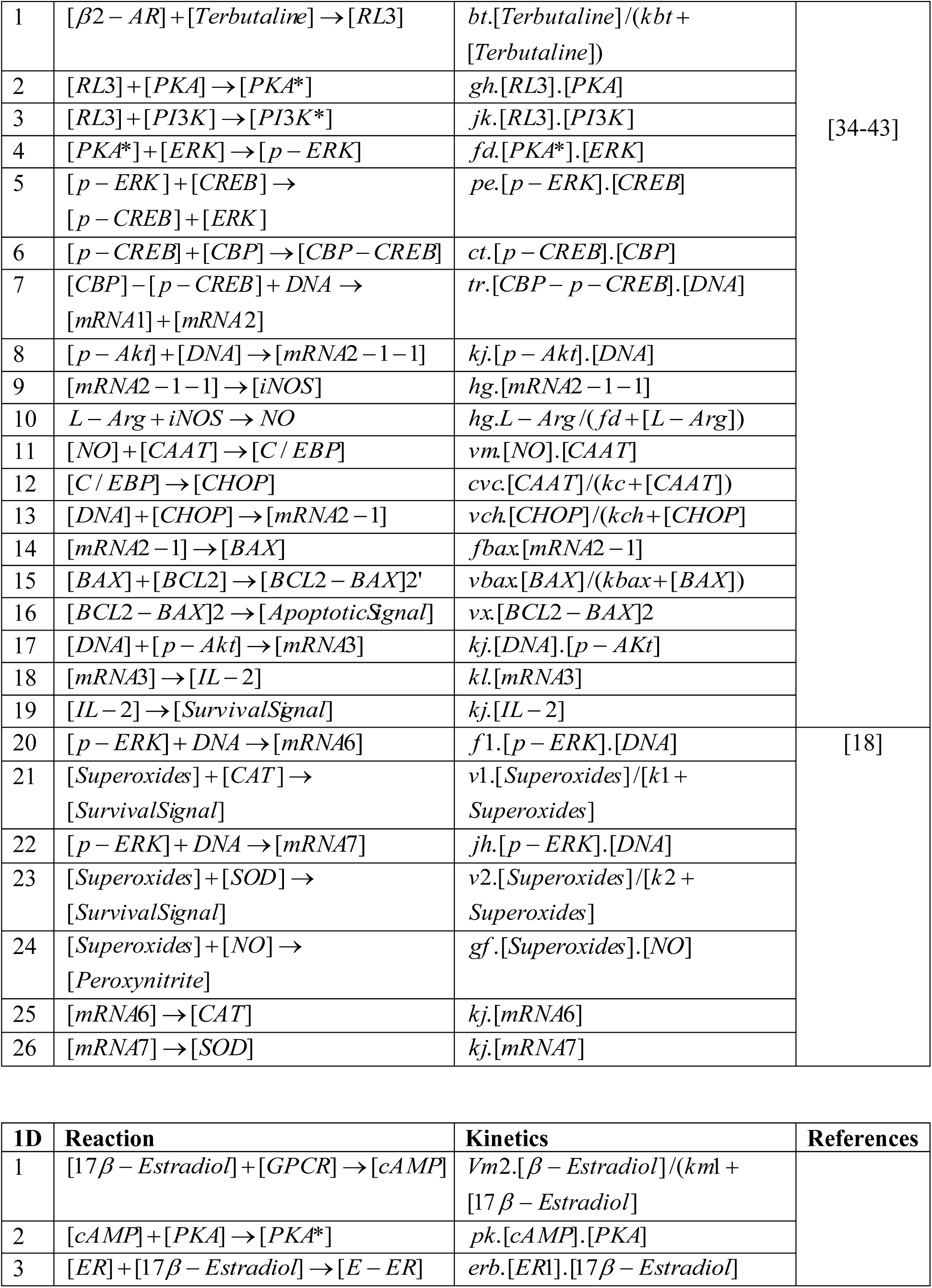
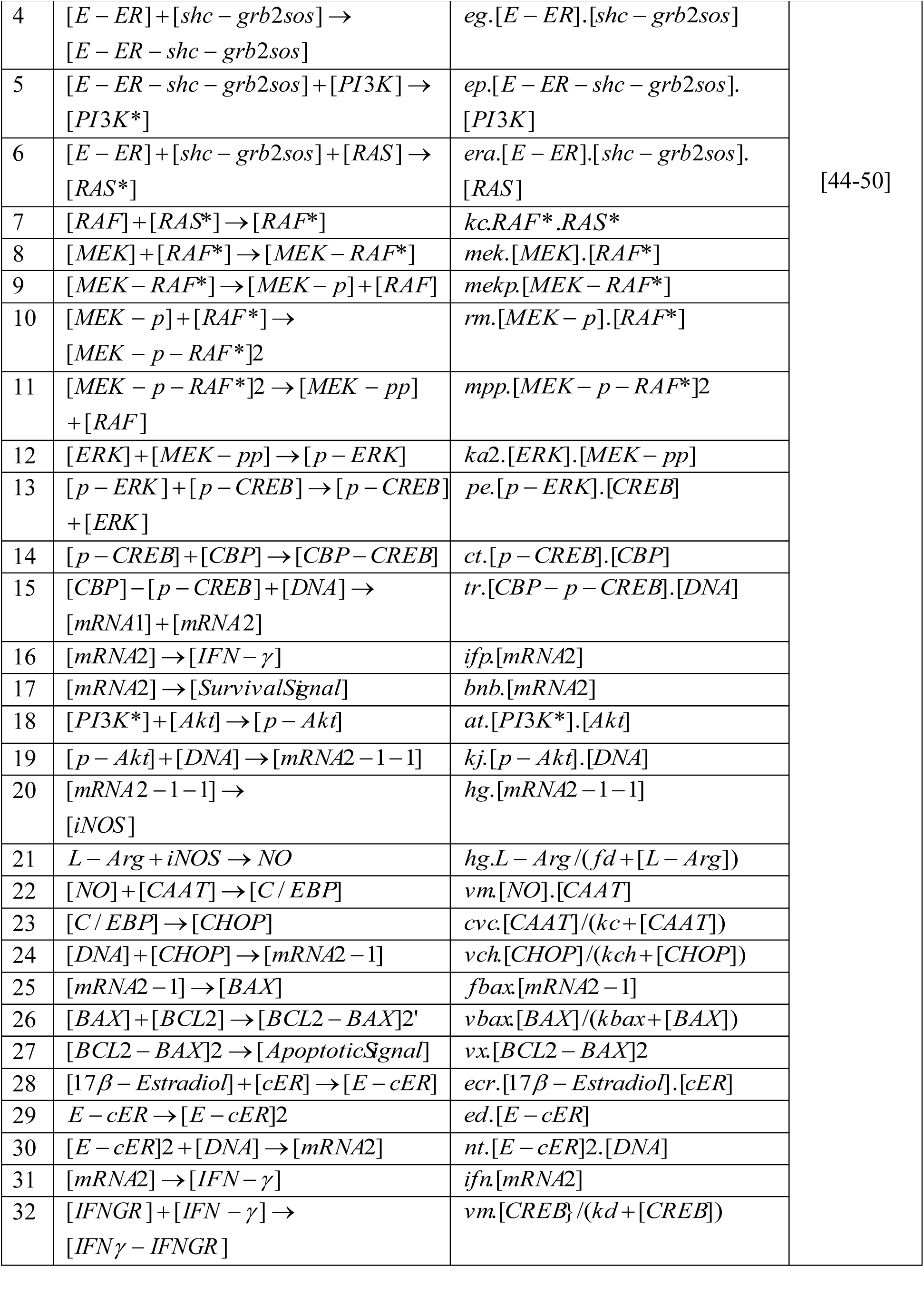
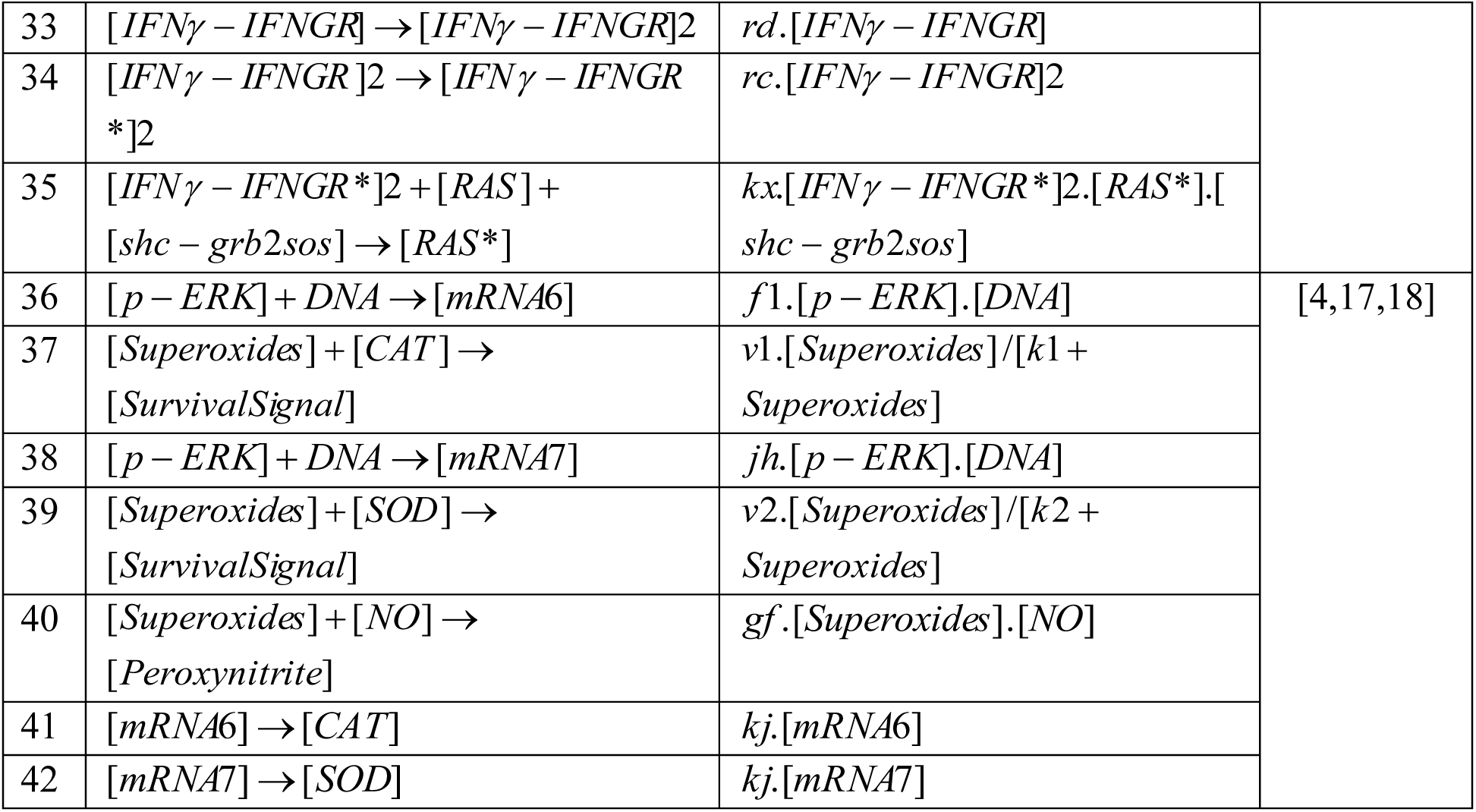
Reaction table and kinetics for α1-adrenoceptor signaling (1A), α2-adrenoceptor signaling (1B), β2-adrenoceptor signaling (1C) and 17β-estradiol signaling through ERs and GPCRs (1D).

**Table 2:** Receptor Binding maxima and Kd values (A), Signaling Molecule concentrations (B) and Ligand concentrations (C) as incorporated in the model.

| 2A | Receptor | Bmax | Kd (nM) | Reference |
| --- | --- | --- | --- | --- |
| 1. | $\alpha 1$-AR | 175.3 (fM/10<sup>6</sup> cells) | 0.65 | [19] |
| 2. | $\alpha 2$-AR | 19.9 (fM/10<sup>6</sup> cells) | 3.7 | [21] |
| 3. | $\beta 2$-AR | 1222 sites/cell | 19.9 | [23,24] |
| 4. | IFN-gR | 708±14 receptors/cell | 0.9±0.2 | [24] |
| 5. | cER | 0.6204 (fM/mg protein) | 5.5 | [25] |
| 6. | nER | (0.136 fm/mcg dna) | - | [25] |

| 2B | Signaling Moelcules | Concentration (nM) | Reference |
| --- | --- | --- | --- |
| 1. | ERK | 251 | [17] |
| 2. | CREB | 245.6 |  |
| 3. | Akt | 219 |  |
| 4. | IFN-g | 560 |  |

| 2C | Signaling Molecules | Concentration | Reference |
| --- | --- | --- | --- |
| 1. | Phenylephrine | $10^{-6}$ M | [17, 18] |
| 2. | Clonidine | $10^{-6}$ M | |
| 3. | Terbutaline | $10^{-6}$ M | |
| 4. | Estrogen | $10^{-8}$ M | |

#### 2.1.1 Model

The model was built to show an interconnected network of four signaling pathways on the basis of in vitro studies conducted in our laboratory including:

1. α1-adrenoceptor signaling [17]; (Table 1A, Figure 1A) Construction of the α1-adrenoceptor signaling pathway was based on the data from our lab and others that showed that stimulating the α1-AR in splenocytes with 10^−6^ M and 10^−9^ M specific agonist phenylephrine did not alter splenocyte proliferation but enhanced survival signaling molecules including p-ERK and p-CREB [17]. In order to simulate the receptor-ligand reactions in real time, the receptor ligand interactions were defined for α1-AR as shown in table 2A [19]. The concentrations of ERK and CREB used in the model were based on the data from the published in vitro study determined using ELISA [17]. Downstream signaling cascades were assumed to follow the law of mass action. Molecules for which concentrations were not known such as PKA and CBP were assigned arbitrary values (257 nM) that did not impose a constraint on the system.
2. α2-adrenoceptor signaling [17]; (Table 1B, Figure 1A): α2-AR signaling was constructed based on the data obtained from our study showing that incubation of α2-AR specific agonist Clonidine (10^−9^ M-10^−6^ M) enhanced Akt signaling cascade [17]. Although our study showed a decrease in cellular proliferation in response to clonidine, there was no reversal upon co-treatment with specific antagonist idazoxan. It is possible that α2-AR stimulation is necessary for the sustenance of nerve fibres through activation of Bcl-2 downstream to Akt signaling [20]. The receptor ligand interactions for α2-AR are defined in table 2A [21]. The concentration for Akt was obtained from the in vitro study using ELISA [17]. Downstream signaling cascades were assumed to follow the law of mass action. Molecules for which concentrations were not known such as BCL-2 were assigned arbitrary values (300 nM) that did not impose a constraint on the system.
3. β2-adrenoceptor signaling [18]; (Table 1C, Figure 1A): Construction of the β2-adrenoceptor signaling was based on the data obtained from the published in vitro study [18]. Stimulation of splenocytes using the β2-adrenoceptor agonist terbutaline significantly enhanced p-ERK and p-CREB expression, activity of SOD and CAT through PKA. Terbutaline treatment also enhanced IL-2 expression through p-Akt. p-Akt expression also enhances the activity of iNOS leading to an increase in NO expression. NO can react with superoxides and form peroxynitrites that may accumulate and lead to apoptosis, or trigger CAAT, CEBP, CHOP, BCL-2, leading to dimerization of BCL-2-Bax complex leading to apoptosis [22]. The receptor ligand interactions for β2-AR and IFN-γ are defined in table 2A [23,24]. The concentration for ERK, CREB, Akt, NO, IFN-γ, IL-2 were obtained from the in vitro study using ELISA [17]. Downstream signaling cascades were assumed to follow the law of mass action. Molecules for which concentrations were not known such as PI3K, PKA, CBP (257 nM), SOD, CAT, iNOS (0 units), CAAT, CEBP, CHOP, BAX, BCL-2, O2-(0 units) were assigned arbitrary initial values that did not impose a constraint on the system.
4. 17β-estradiol signaling through ERs and GPCRs [4]; (Table 1D, Figure 1B) and their respective kinetic parameters outlined in Table 2. Estrogen signaling was constructed based on the data from the in vitro study published from our lab [4]. Splenocytes stimulated with estrogen showed enhanced expression of ERK and CREB signals through PKA and cAMP by estrogen binding to GPCRs, leading to the expression of IFN-γ. IFN-γ can bind to IFN-γ receptor, activate PI3K, Ras, Raf, MEK, ERK and CREB cascades leading to survival signals. Estrogen can bind to estrogen receptors, stimulate Shc, Grb-2, Sos, activate PI3K, Akt, iNOS leading to NO expression, activation of CAAT, CEBP, CHOP and induce pro-apoptotic signals through Bcl-2/Bax cascades. NO can also bind with superoxides leading to the production of peroxynitrites that serve as an apoptotic signal. On the other hand, Estrogen also promotes the expression of SOD and CAT through p-ERK leading to scavenging of the excess superoxides leading to survival signals [4]. Estrogen can also bind to nuclear receptors and lead to the production of IFN-γ. The receptor ligand interactions for estrogen receptors, IFN-γ receptors are defined in table 2A [24,25]. The concentration for ERK, CREB, Akt, NO, IFN-γ were obtained from the in vitro study using ELISA [17]. Downstream signaling cascades were assumed to follow the law of mass action. Molecules for which concentrations were not known such as PI3K, PKA, CBP (257 nM), SOD, CAT, iNOS (0 units), CAAT, CEBP, CHOP, BAX, BCL-2, O2- (0 units) were assigned arbitrary initial values that did not impose a constraint on the system. Enzyme kinetics for CAT, SOD and iNOS were defined using the Henri-Michaelis-Menten equation.

### 2.2 Simulations

Simulations were performed using Ode15s solvers for 24 hours and 72 hours depending upon the signals studied. The Mcode is included in the supplementary evidence. The model was tested for various combinations as follows and the figures are shown in supplementary evidence:

**Figure 1.**
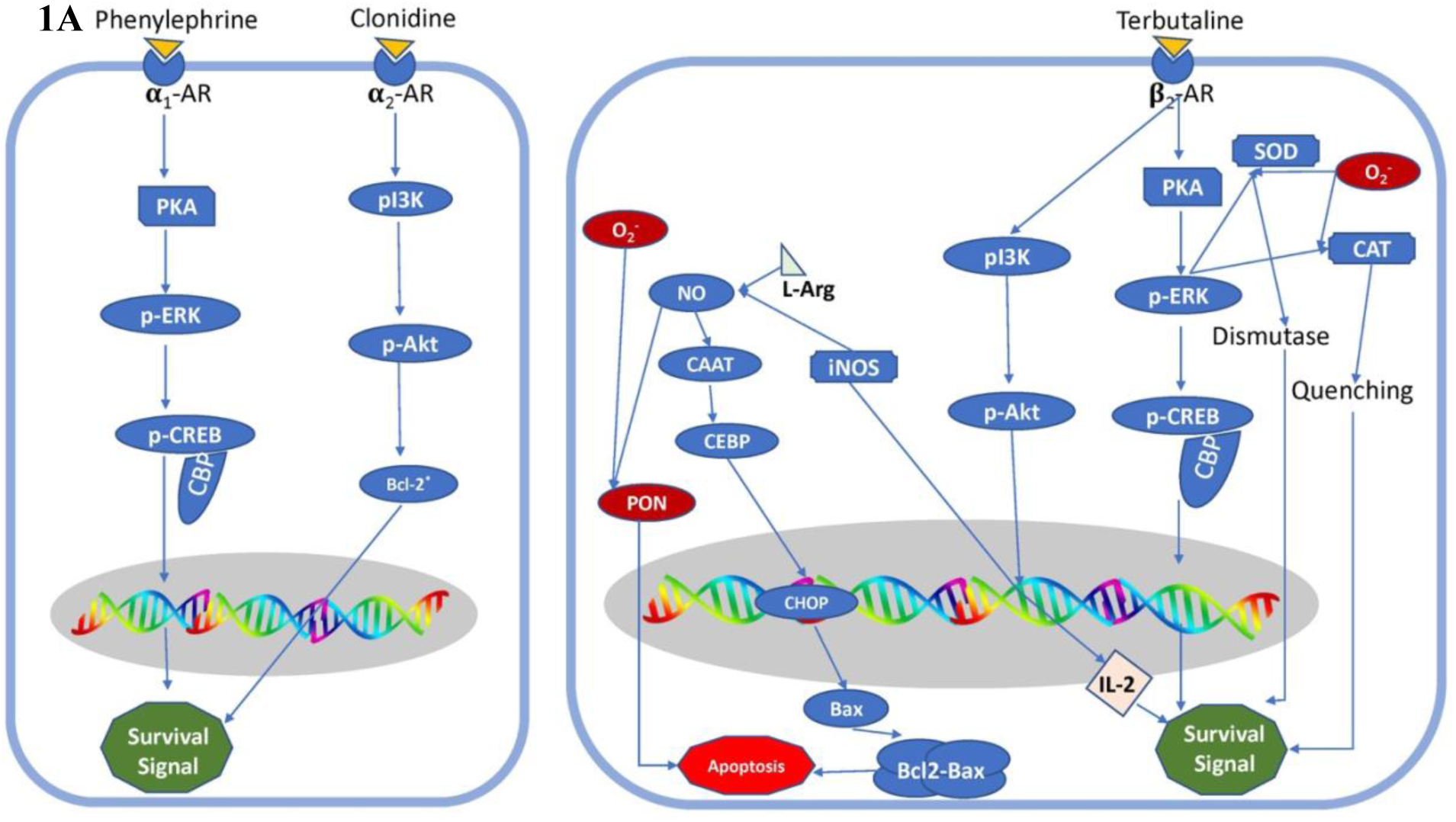
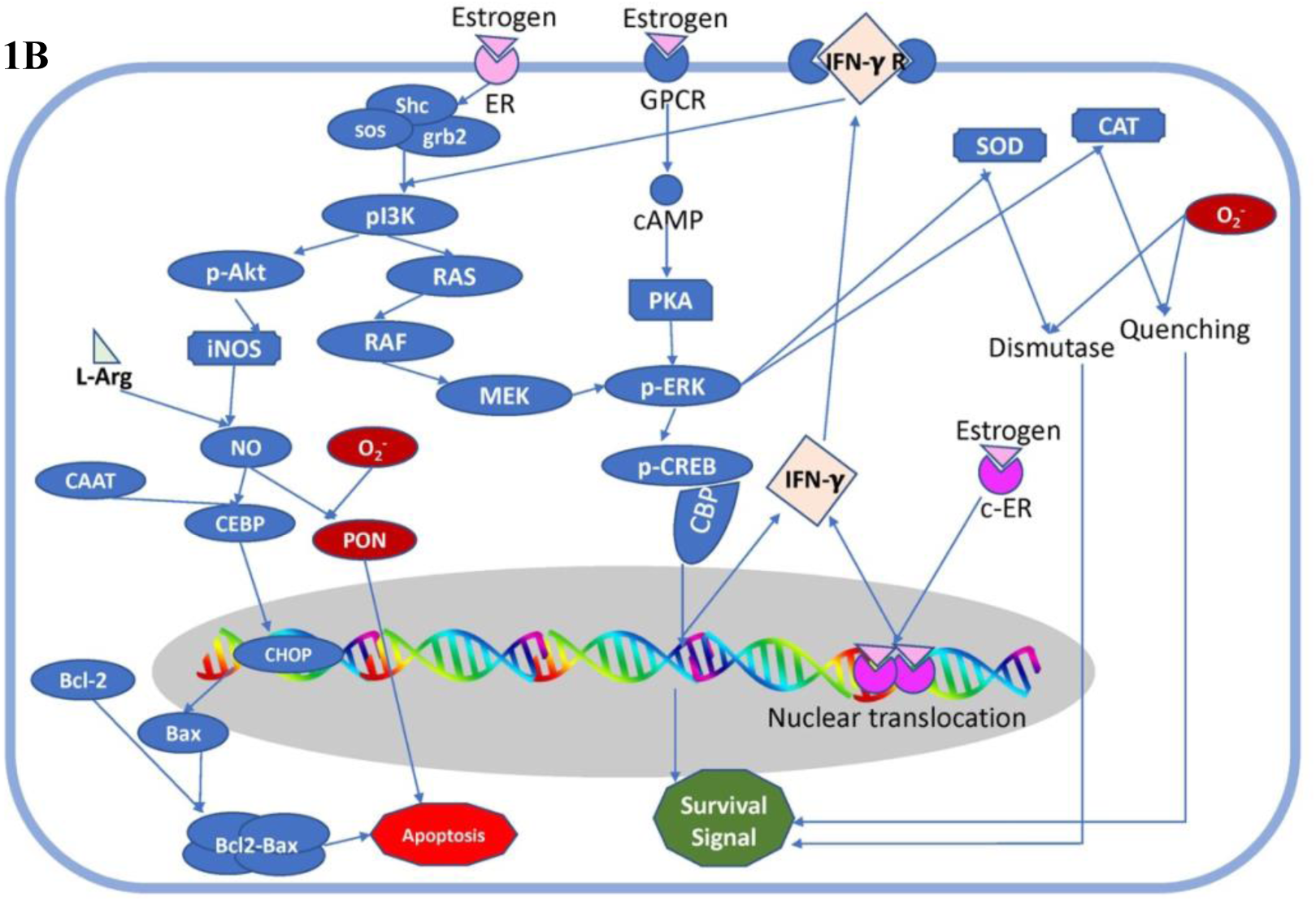
In silico model of adrenergic [1A] and estrogen [1B] signaling cascades in lymphocytes: The cross-talk between the adrenergic and estrogen-mediated signals result in specific immunomodulatory effects depending upon the estrogen concentration. The layout of the signaling pathway is elucidated from receptors to transduction into the cytosol and nuclear translocation of signaling molecules leading to specific outcomes.

A. Estrogen Signaling: (Supplementary Figures 1-4)
  a. Simulation Conditions:
    i. Terbutaline=0 M;
    ii. Phenylephrine=0 M;
    iii. Clonidine=0 M;
    iv. E_2_= 10^−3^ M, 10^−6^ M, 10^−9^ M
    v. Time : 24 hours, 72 hours
B. Adrenergic Signaling:
  a. Simulation Condition-1: Terbutaline Signaling (Supplementary Figures 5-8)
    i. Terbutaline= 10^−3^ M, 10^−6^ M, 10^−9^ M;
    ii. Phenylephrine= 10-6 M;
    iii. Clonidine= 10^−6^ M;
    iv. E_2_= 0 M
    v. Time : 24 hours, 72 hours
  b. Simulation Condition-2: Phenylephrine Signaling (Supplementary Figures 9-12)
    i. Terbutaline= 10^−6^ M;
    ii. Phenylephrine= 10^−3^ M, 10^−6^ M, 10^−9^ M;
    iii. Clonidine= 10^−6^ M;
    iv. E_2_= 0 M
    v. Time : 24 hours, 72 hours
  c. Simulation Condition-3: Clonidine Signaling (Supplementary Figures 13-16)
    i. Terbutaline= 10^−6^ M;
    ii. Phenylephrine= 10^−6^ M;
    iii. Clonidine= 10^−3^ M, 10^−6^ M, 10^−9^ M;
    iv. E_2_= 0 M
    v. Time : 24 hours, 72 hours
C. Adrenergic+Estrogen Signaling
  a. Simulation Condition-1: Terbutaline+Estrogen Signaling (Supplementary Figures 17-20)
    i. Terbutaline= 10^−3^ M, 10^−6^ M, 10^−9^ M;
    ii. Phenylephrine= 10^−6^ M;
    iii. Clonidine= 10^−6^ M;
    iv. E_2_= 10^−8^ M
    v. Time : 24 hours, 72 hours
  b. Simulation Condition-2: Phenylephrine+Estrogen Signaling (Supplementary Figures 21-24)
    i. Terbutaline= 10^−6^ M;
    ii. Phenylephrine= 10^−3^ M, 10^−6^ M, 10^−9^ M;
    iii. Clonidine= 10^−6^ M;
    iv. E_2_= 10^−8^ M
    v. Time : 24 hours, 72 hours
  c. Simulation Condition-3: Clonidine+Estrogen Signaling (Supplementary Figures 24-28)
    i. Terbutaline= 10^−6^ M;
    ii. Phenylephrine= 10^−6^ M;
    iii. Clonidine= 10^−3^ M, 10^−6^ M, 10^−9^ M;
    iv. E_2_= 10^−8^ M
    v. Time : 24 hours, 72 hours

For signaling molecules including p-ERK, p-Akt, p-CREB, and cAMP, cytokines including IL-2 and IFN-γ, antioxidant enzymes (SOD and CAT) and superoxides/peroxynitrites, the simulations were performed up to 24 hours similar to the in vitro study. Finally, for survival vs. apoptotic signals, the simulations were carried out for 72 hours. Since all the adrenergic receptors are activated by the same ligand norepinephrine, physiologically, the simulations were run for different doses of one receptor subtype (e.g., terbutaline 10^−3^ M, 10^−6^ M, 10^−9^ M) while allowing for a mid-range activity in the other two (clonidine 10^−6^ M, phenylephrine 10^−6^ M) in the presence (10^−8^ M) and absence (0 M) of estrogen. Key regulatory molecules were scanned, and their sensitivities were analyzed at 24 hours for each of the ligands used (Supplementary Figures 1D-28D; M Code attached as Supplementary Index-2). Parameter sensitivity analysis was run for the initial concentrations of all the parameters included in the study (Table 3).

**Table 3:**
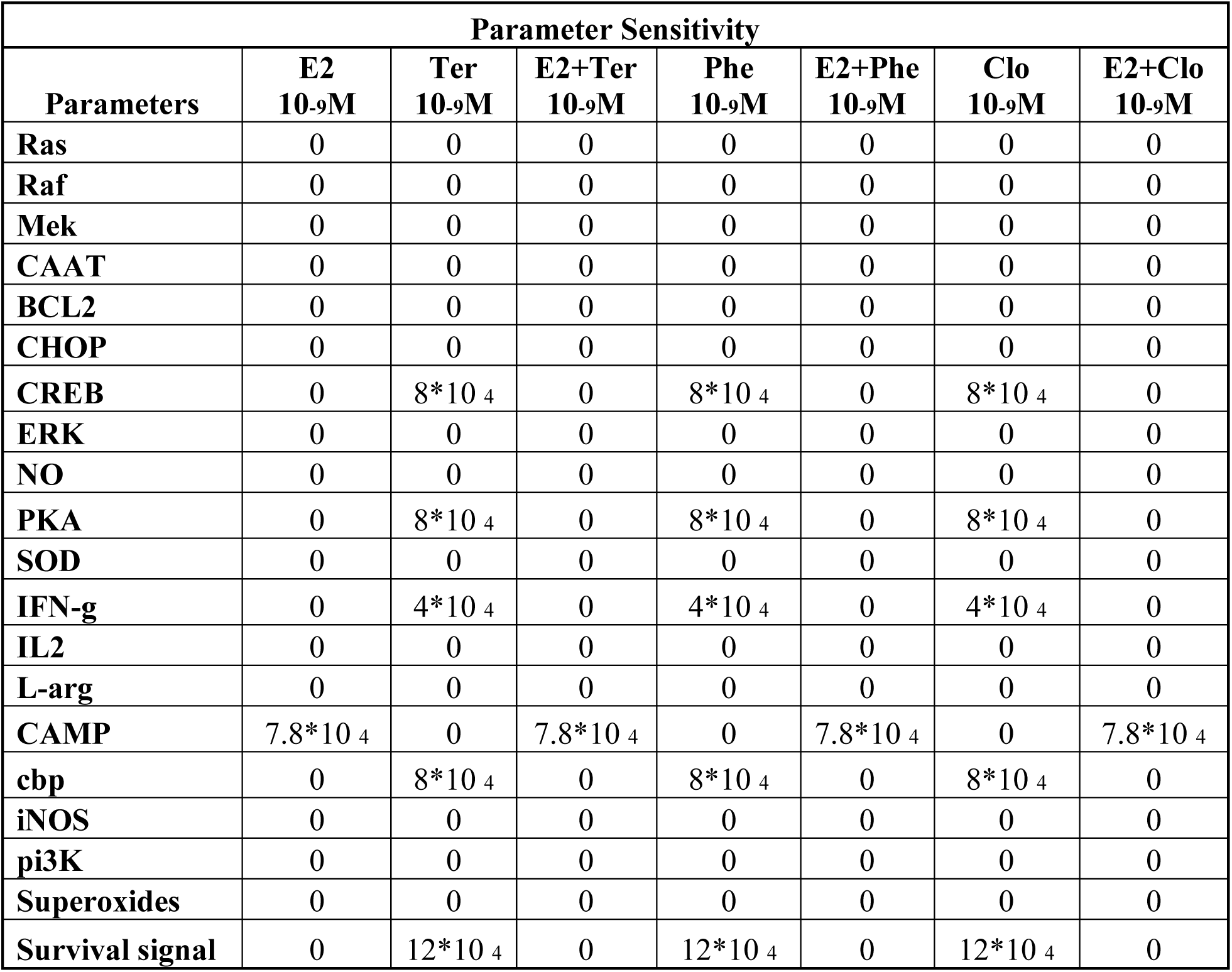
Parameter sensitivity scan in fM for 24 hours for Estrogen (E^2^ 10^−9^M), and adrenergic agonists, Terbutaline (Ter 10^−9^ M), Phenylephrine (Phe 10^−9^M), Clonidine (Clo 10^−9^M), alone and in combination with estrogen.

## 3.0 Results and Discussion

Immune functions of innate, humoral or cell-mediated origin are modulated by sympathetic signals and circulating hormone levels during health and disease in a dose and receptor-type-dependent manner [1, 10, 12, 51]. *In vitro* studies from our laboratory have shown that adrenergic stimulation through α1-, α2- or β2-ARs using specific agonists non-specifically inhibit lymphoproliferation through distinct signaling pathways: α1-ARs mediated immunosuppressive effects by inhibiting IFN-γ production, α2-AR activated the NF-κB, p-Akt and NO pathways and β2-AR activation involved IL-6, NO and NF-κB signaling cascades mediating immunosuppression [17, 18]. On the other hand, treatment of lymphocytes with 17β-estradiol enhanced proliferation in a dose and receptor subtype specific manner through p-ERK, p-CREB and p-Akt involving IFN-γ and compensatory mechanisms including antioxidant enzymes [4, 52]. Our studies have shown that co-treatment of lymphocytes with 17β-estradiol and adrenergic agonists lead to 17β-estradiol-mediated over-ride of adrenergic immunosuppression in a dose-dependent, AR-subtype independent manner [17, 18].

In the present study, we have modelled these signaling pathways *in silico* using the MATLAB Simbiology toolbox. Due to difficulties in conducting the experiments using NE, the *in vitro* studies were conducted using receptor-specific agonists and antagonists. While these experiments are highly beneficial in elucidating the dose-dependent effects on specific adrenoceptor sub-types, they cannot be considered as absolute indicators of NE-mediated effects. This is because when NE is added to the cells, depending on its relative affinity for the individual receptor subtypes, it may bind proportionally to all the ARs triggering a wide variety of down-stream signals. The cumulative effects of these signals are likely to be different from the outcomes of receptor-subtype-specific signaling fates. In order to overcome this problem, the computational model was constructed on the basis of the data obtained from *in vitro* studies and all three Ars studied were simultaneously activated to study the outcome. Based on previous findings, an attempt was made to incorporate cross-talks between these pathways by using common signaling molecule pools (Fig. 1).

As expected, the simulation of lymphocytes with adrenergic agonists, phenylephrine, clonidine, and terbutaline enhanced p-CREB and p-ERK expression (Fig. 3). 17β-estradiol behaved similarly in the presence and absence of adrenergic agonists enhancing cAMP, p-ERK, and p-CREB expression suggesting that activation of cAMP may be crucial to 17β-estradiol-mediated over-ride of adrenergic immunosuppression (Fig. 2 and 4).

**Figure 2.**
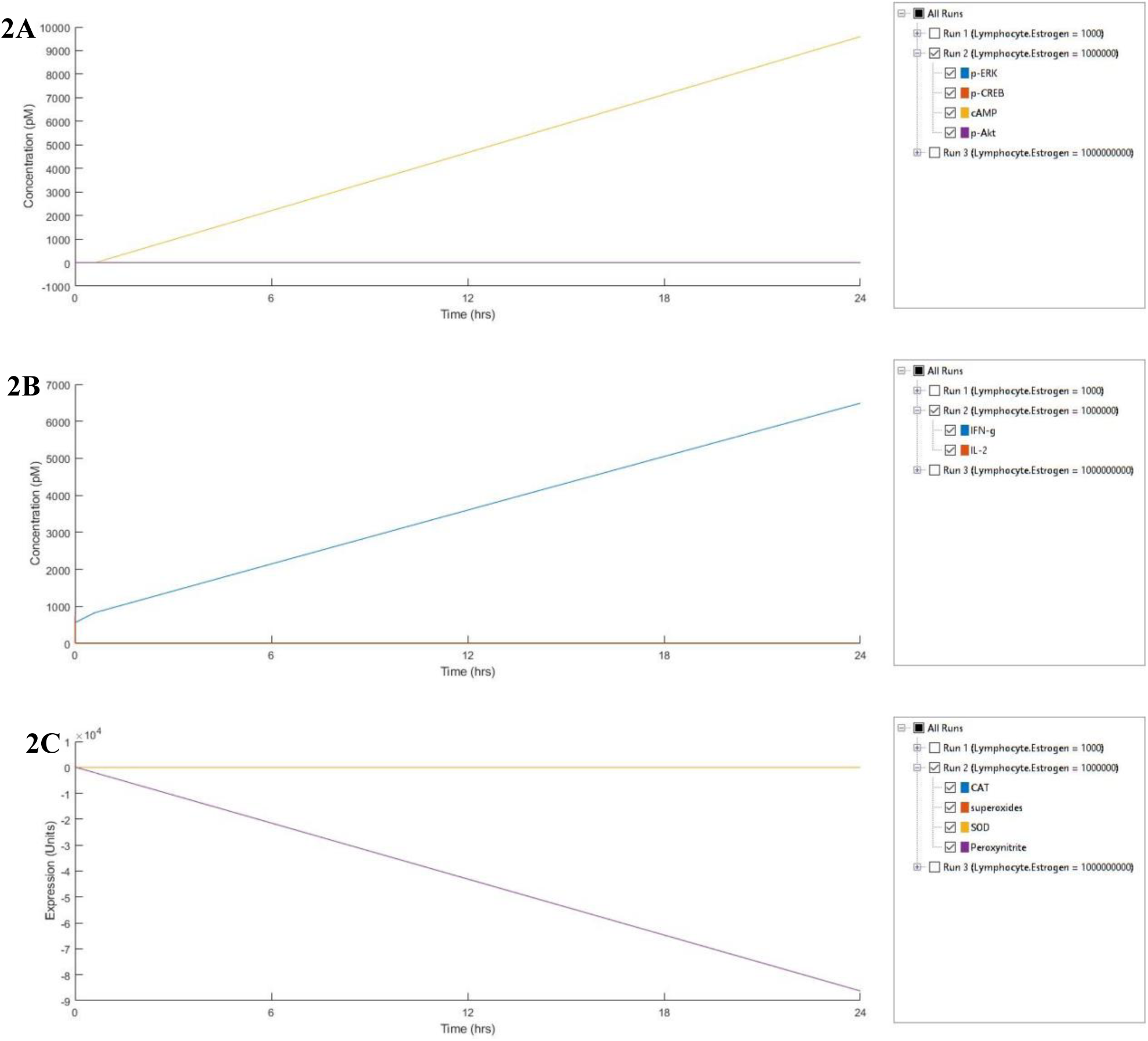
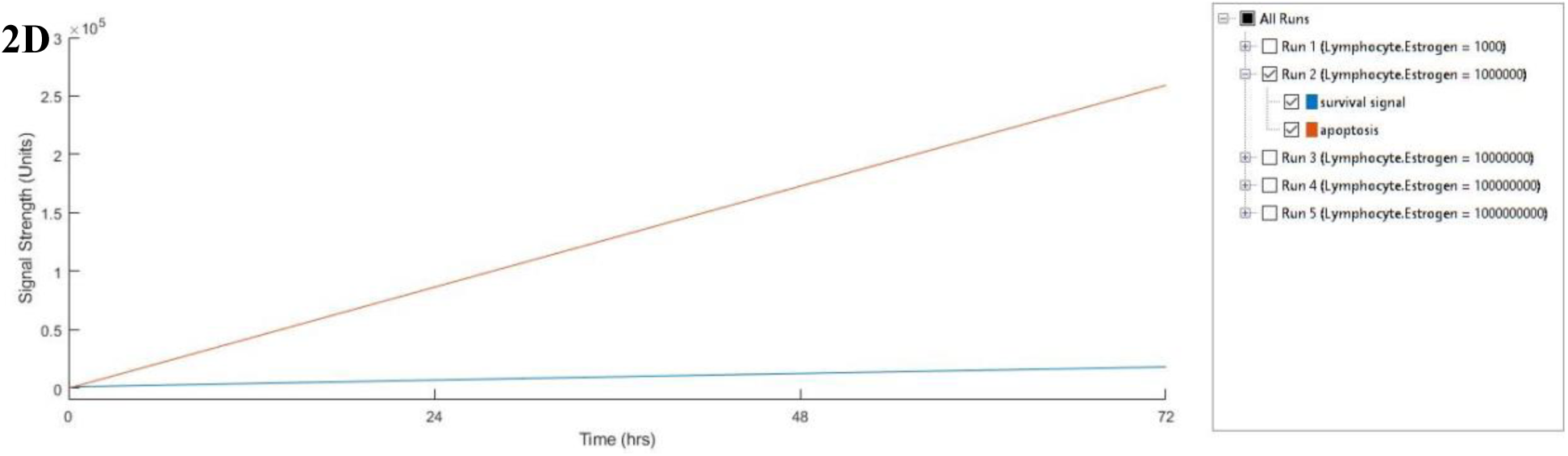
Expression of molecular markers (2A), cytokines (2B), antioxidant enzymes (2C) and survival/apoptosis signals (2D) by estrogen: Simulation of lymphocytes treated with 10^−6^ M Estrogen for 24 hours.

**Figure 3.**
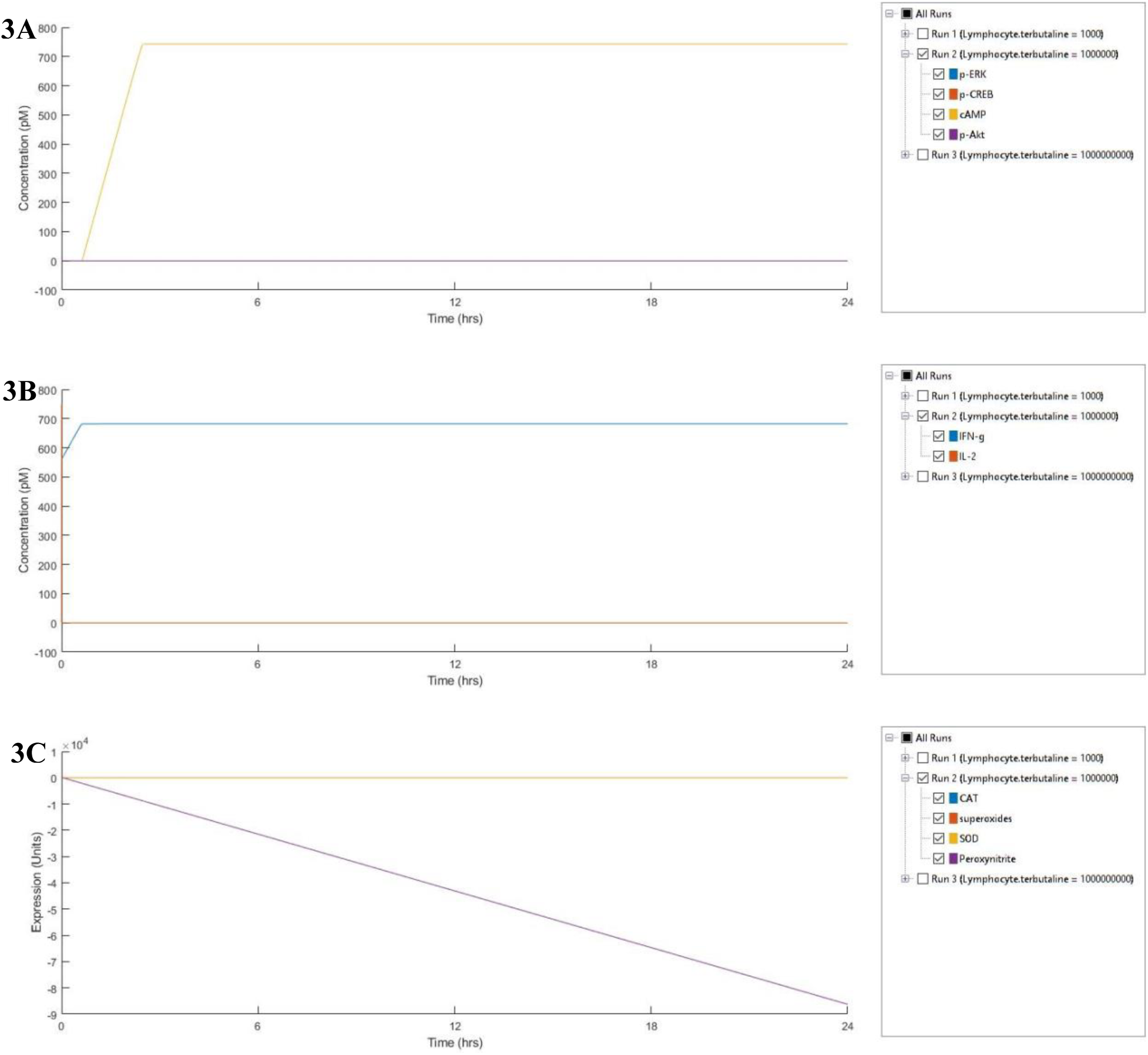
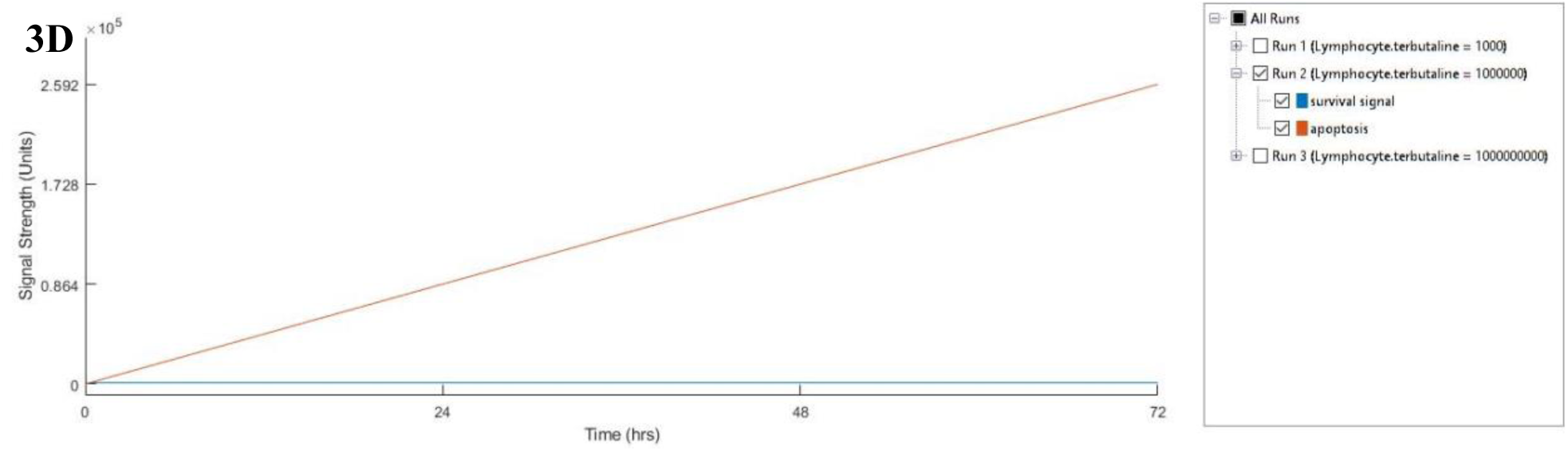
Expression of molecular markers (3A), cytokines (3B), antioxidant enzymes (3C) and survival/apoptosis signals (3D) by terbutaline, phenylephrine and clonidine: Simulation of lymphocytes treated with terbutaline (10^−6^ M), phenylephrine (10^−6^ M) and clonidine (10^−6^ M) for 24 hours.

**Figure 4.**
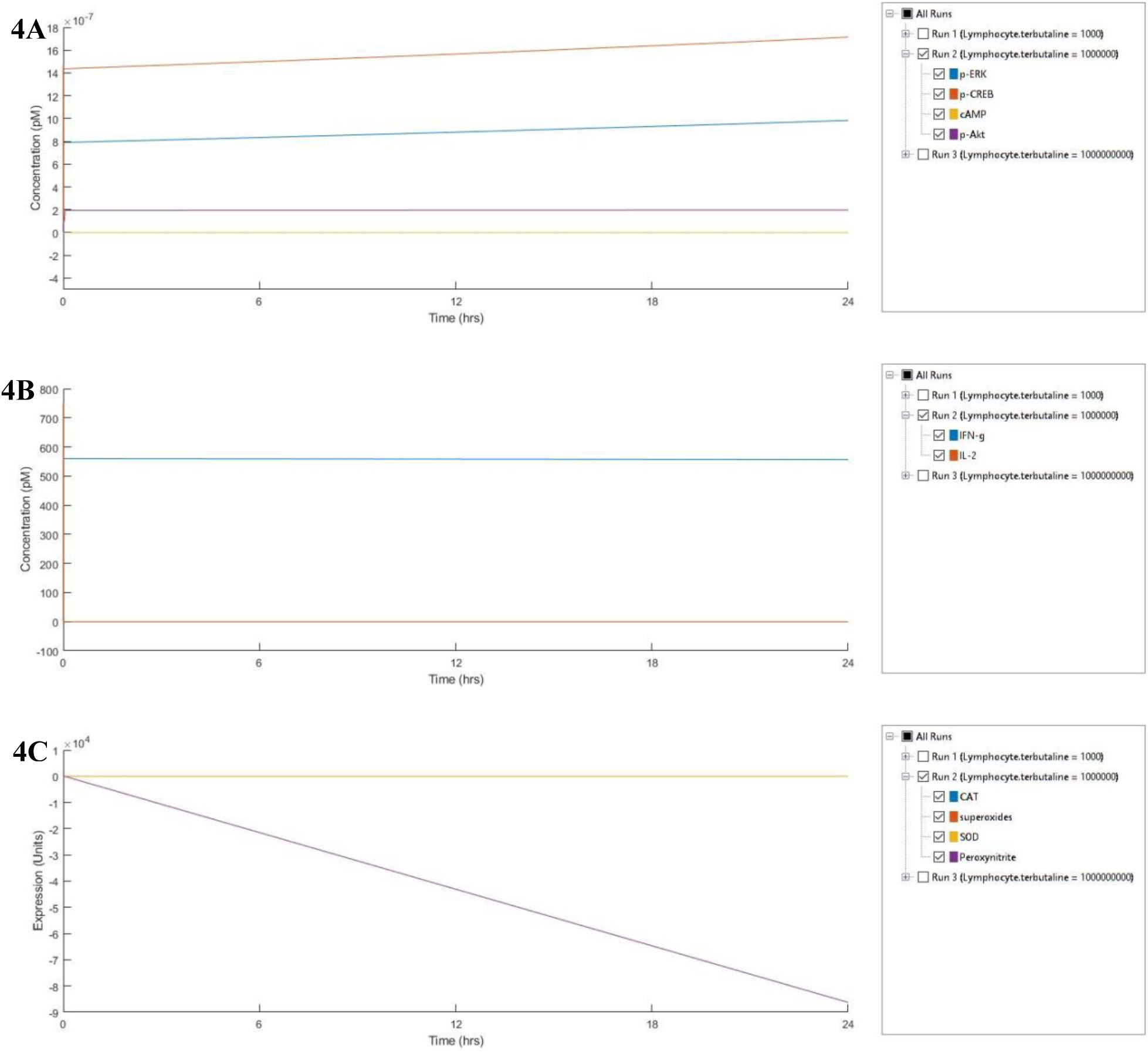
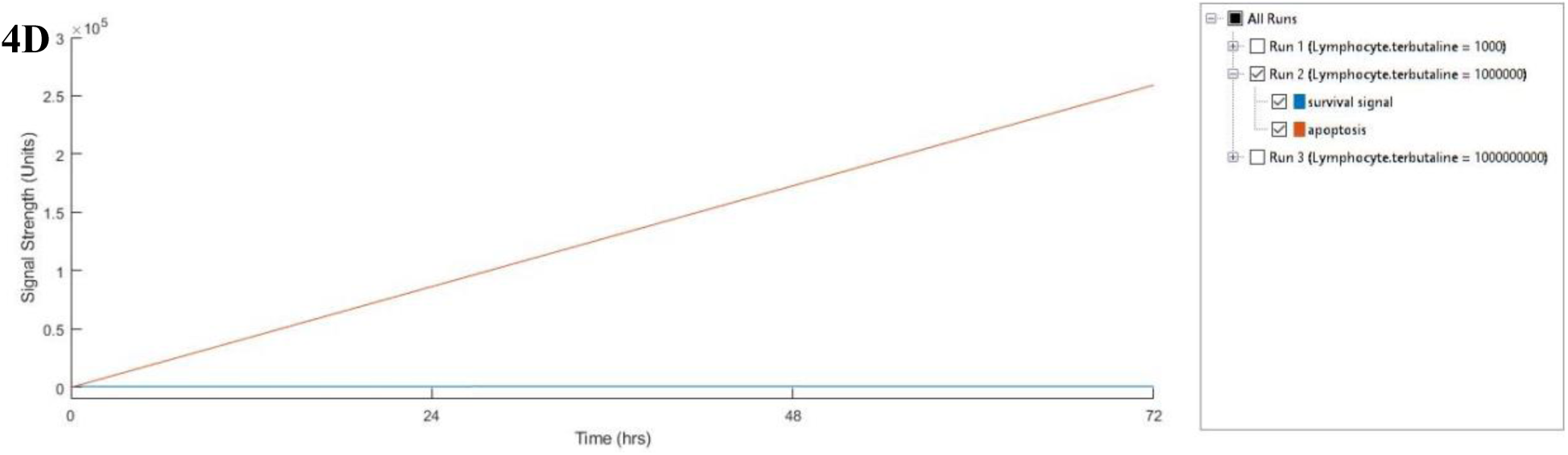
Expression of molecular markers (4A), cytokines (4B), antioxidant enzymes (4C) and survival/apoptosis signals (4D) by terbutaline, phenylephrine and clonidine in the presence of estrogen: Simulation of lymphocytes treated with terbutaline (10^−6^ M), phenylephrine (10^−6^ M) and clonidine (10^−6^ M) in the presence of estrogen (10^−6^ M) for 24 hours.

A significant role is played by compensatory mechanisms such as antioxidant enzyme apart from signaling molecules, to influence the proliferative/apoptotic milieu of a cell by balancing free radical load including superoxides and peroxynitrites to name a few. These free radicals released as by-products following immune responses or during neurotransmitter release in secondary lymphoid organs may accumulate over the years and contribute to denervation of sympathetic noradrenergic fibers leading to impaired neuroendocrine-immune homeostasis setting the stage for age-associated diseases [7]. Studies from our lab and others have documented ER-subtype dependent increase in SOD and GPx activities in lymphocytes stimulated with estrogen [4, 53-58,]. Adrenergic stimulation also enhanced SOD and catalase activities *in vitro* which were suppressed upon coincubation with 17β-estradiol indicating cross-talk between the two pathways leading to diminutive effects [18]. Along these lines, previous studies have implicated the involvement of PKA and ERK pathways, which when inhibited, reversed terbutaline-mediated increase in SOD and CAT activities [4, 17,18]. It is possible that these signaling molecules play a regulatory role in the expression and activity of these enzymes, thereby, exerting immunomodulatory effects [59,60]. In our model, adrenergic activation, 17β-estradiol treatment, and their co-treatment decrease superoxide formation but enhance peroxynitrites, although 17β-estradiol treatment or co-treatment alone was accompanied by an induction of antioxidant enzyme activities (SOD and CAT) (Fig. 2-4).

Cytokines play a crucial role in influencing immune functions thereby affecting survival/apoptosis. Previously we have shown that while activation of α1-ARs decrease IFN-γ and increase IL-2 production, α2- and β2-AR activation did not alter both IFN-γ and IL-2 production [17,18]. However, treatment with 17β-estradiol alone or 17β-estradiol with adrenergic agonists significantly enhance IFN-γ production alone after 24 hours of treatment. We have reported dose-dependent increase in IFN-γ production with 17β-estradiol alone or co-treated with α1- and β2-AR agonists. In agreement with these findings, sensitivity analysis show that stimulation of lymphocytes with α1-AR agonist, phenylephrine, showed increased sensitivity to IFN-γ and p-CREB expression while β2-AR agonist terbutaline showed increased sensitivity to IFN-γ, p-ERK and p-CREB expression (Fig. 3). Interestingly 17β-estradiol signaling showed the highest sensitivity to IFN-γ, p-ERK, p-CREB, and cAMP suggesting a probable cause in the dynamics of 17β-estradiol-mediated signals that predispose it to over-ride adrenergic signals (Fig. 2 and 4).

In order to assess the likely effects of the signals and their cross-talks on proliferation, end-point signals were classified as either pro-apoptotic or survival signals. Plots obtained at the end of 72 hours indicate that while adrenergic stimulation was predominantly pro-apoptotic signals, 17β-estradiol signals were pro-survival (Fig. 2 and 3). Dose-dependent role of 17β-estradiol could be seen by increasing the 17β-estradiol concentration by 10-fold leading to further increase in proliferation signals reversing the balance in favor of proliferation over apoptosis after 72 hours (Fig. 5). This is in accordance with the *in vitro* evidence generated in previous studies where all the adrenergic agonists decreased proliferation while 17β-estradiol played a dose-dependent role in increasing proliferation of lymphocytes.

**Figure 5.**
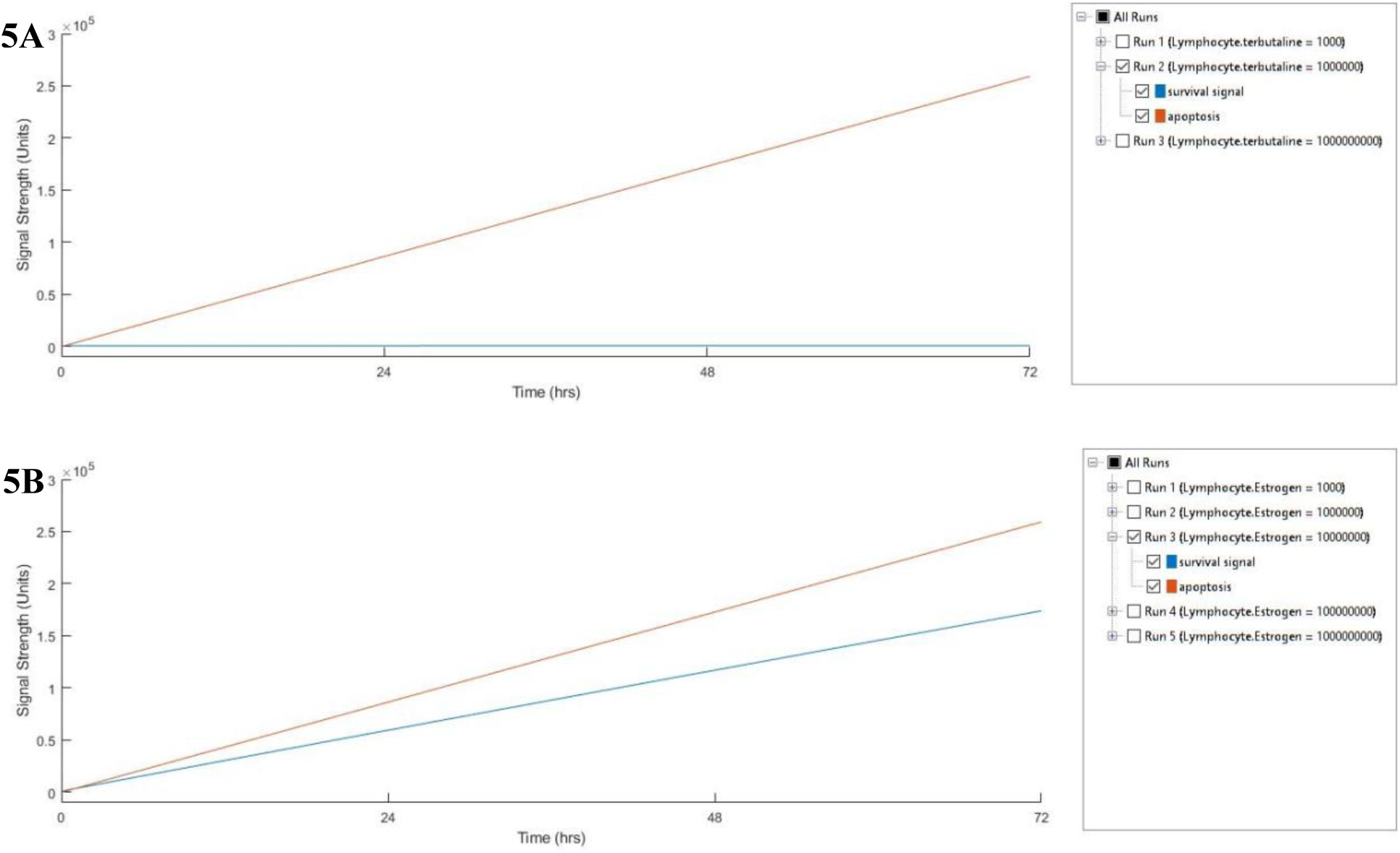
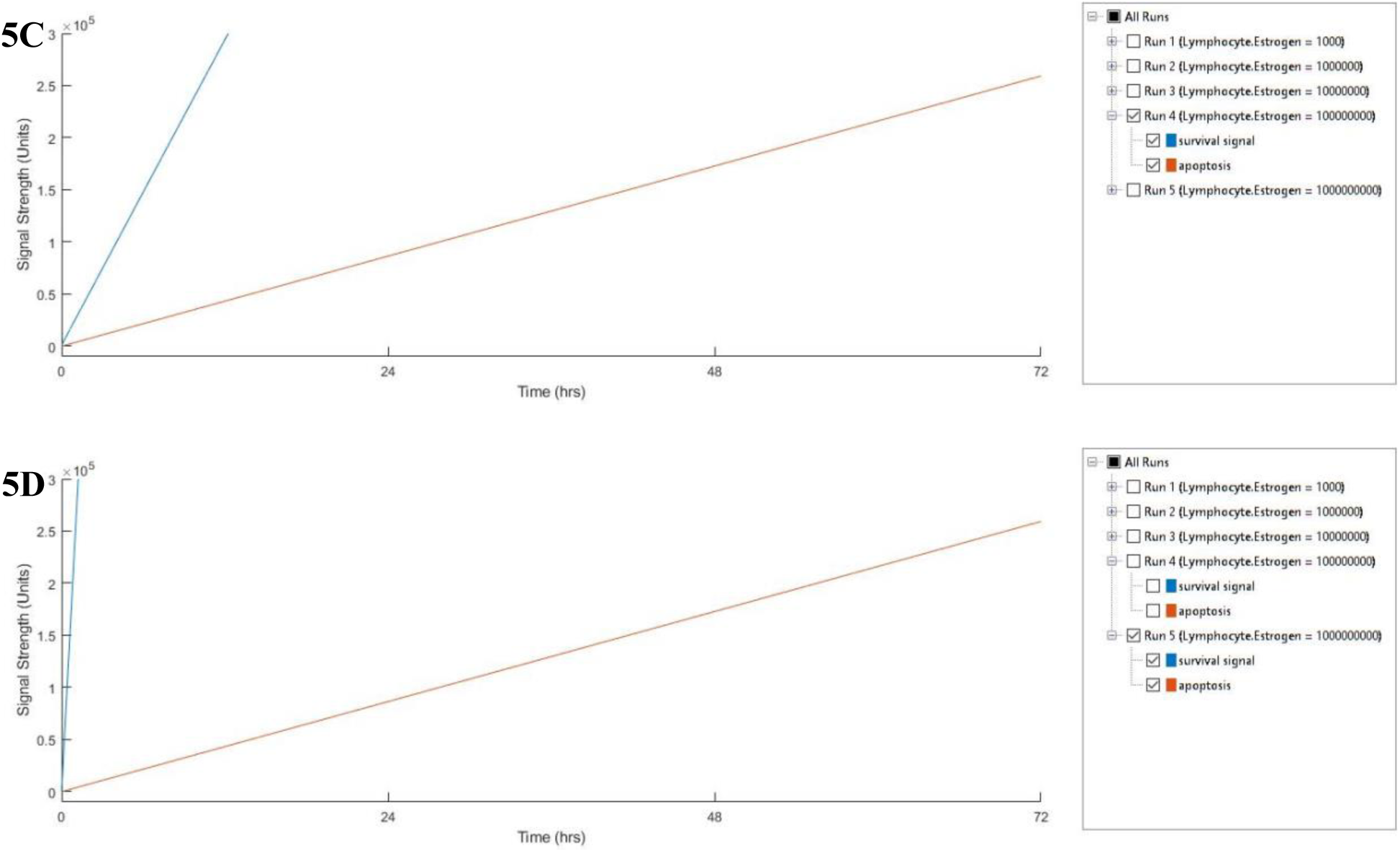
Effects of increasing doses of estrogen on survival/apoptosis signals by terbutaline, phenylephrine and clonidine: Simulation of lymphocytes treated with terbutaline (10^−6^ M), phenylephrine (10^−6^ M) and clonidine (10^−6^ M) in the presence of increasing doses of estrogen (10^−6^ M (5A), 10^−5^ M (5B), 10^−4^ M (5C), 10^−3^ M (5D)) for 72 hours.

Modeling estrogen and adrenergic stimulation-mediated signals will help us to understand variations in the expression of downstream molecular markers with age- and disease-associated variations in kinetic parameters and concentrations and thereby providing an effective tool for understanding the alterations in the cross talk between the pathways in health, aging, and diseases.

## Acknowledgement

Supported by the Department of Science and Technology (F.No. SR/SO/HS-46/2007 and IFA15/LSBM-154), Government of India, New Delhi.

## Supplementary Figures

**Supplementary Figure 1:**
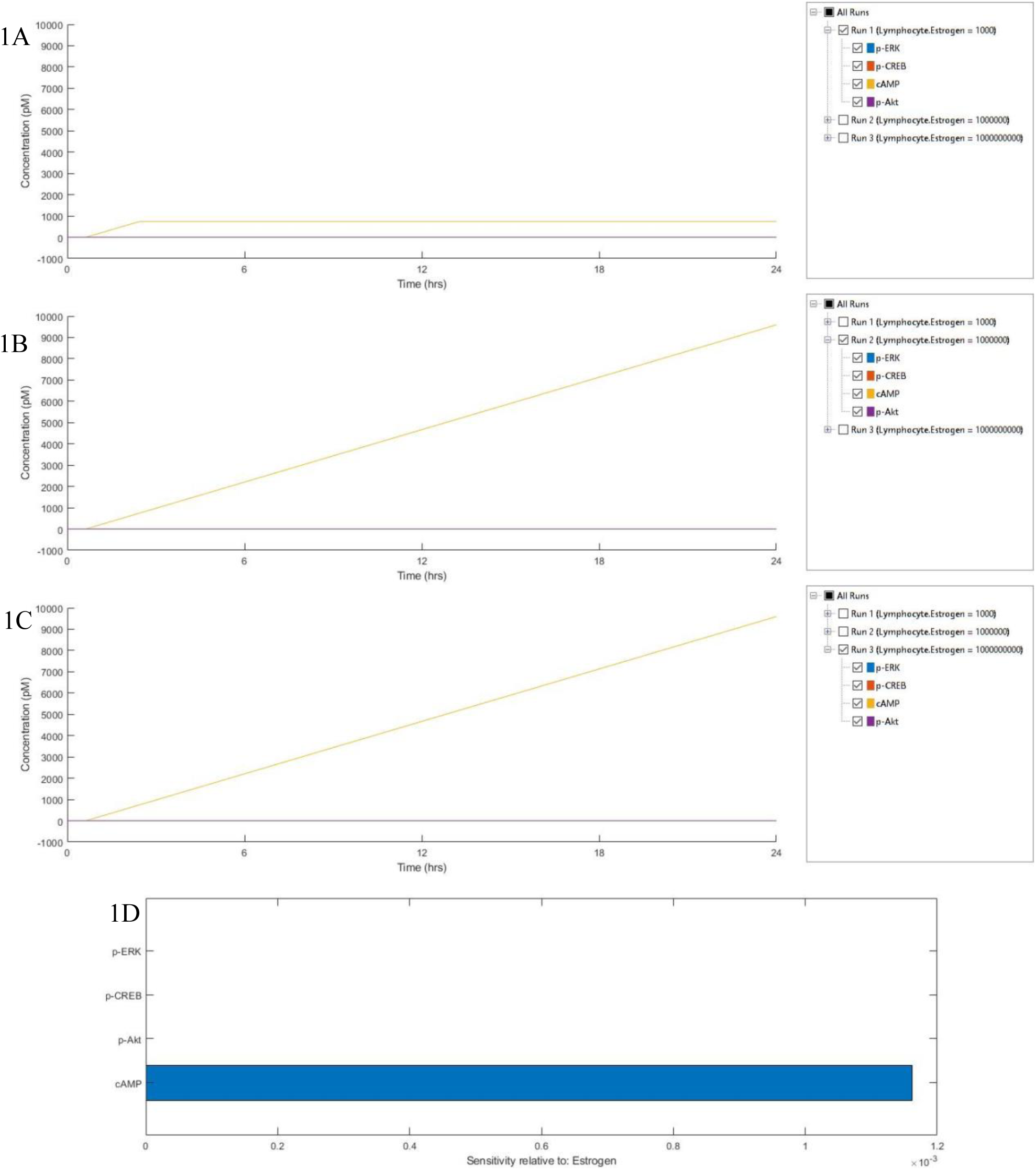
Simulation of Estrogen Signaling: Expression of molecular markers: p-ERK, p-CREB, cAMP and p-Akt upon stimulation with Estrogen 10^−9^M (A), 10^−6^M (B) and 10^−3^M (C) after 24 hours and their sensitivities relative to Estrogen (D).

**Supplementary Figure 2:**
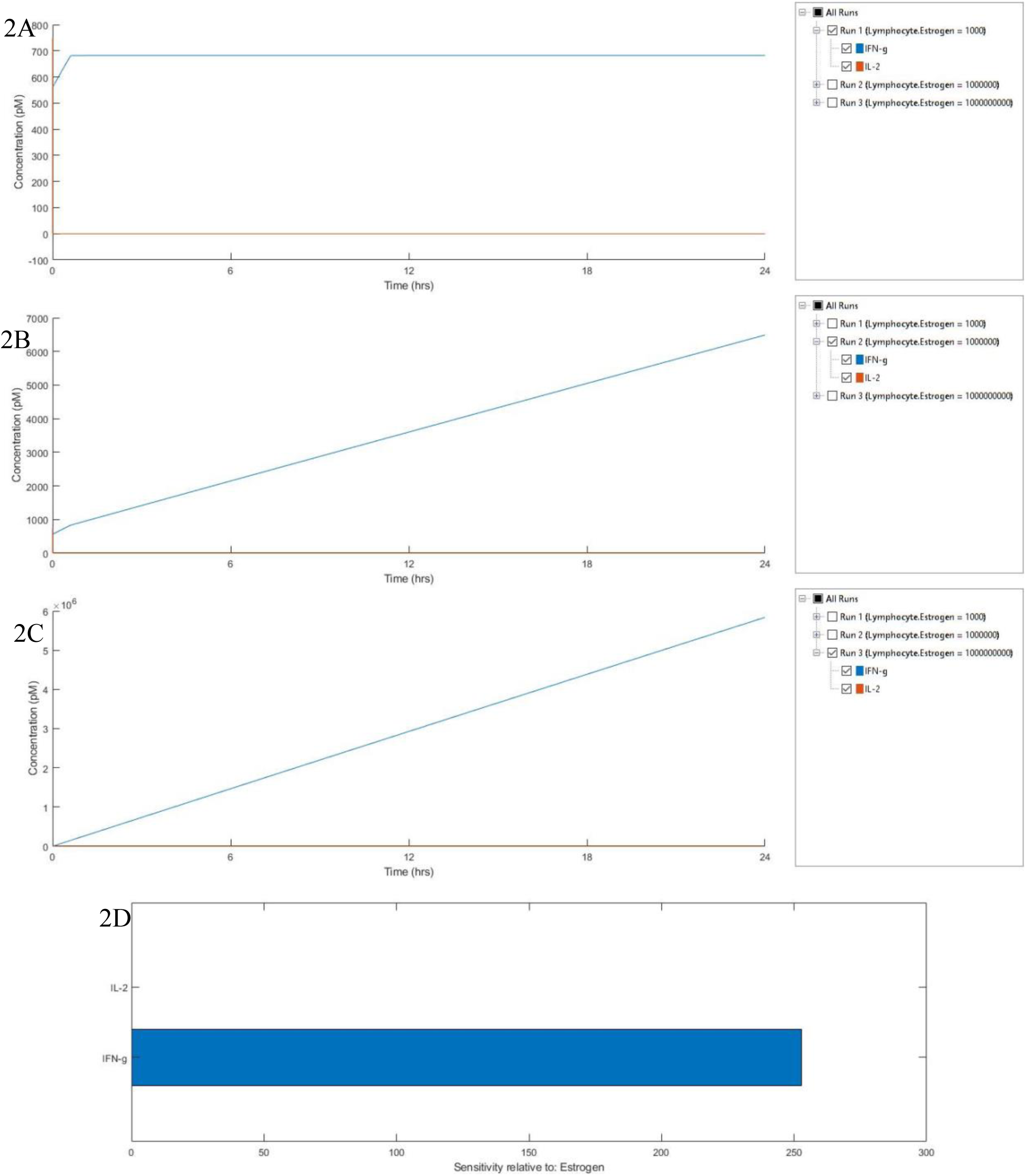
Simulation of Estrogen Signaling: Expression of cytokines: IFN-γ and IL-2 upon stimulation with Estrogen 10^−9^M (A), 10^−6^M (B) and 10^−3^M (C) after 24 hours and their sensitivities relative to Estrogen (D).

**Supplementary Figure 3:**
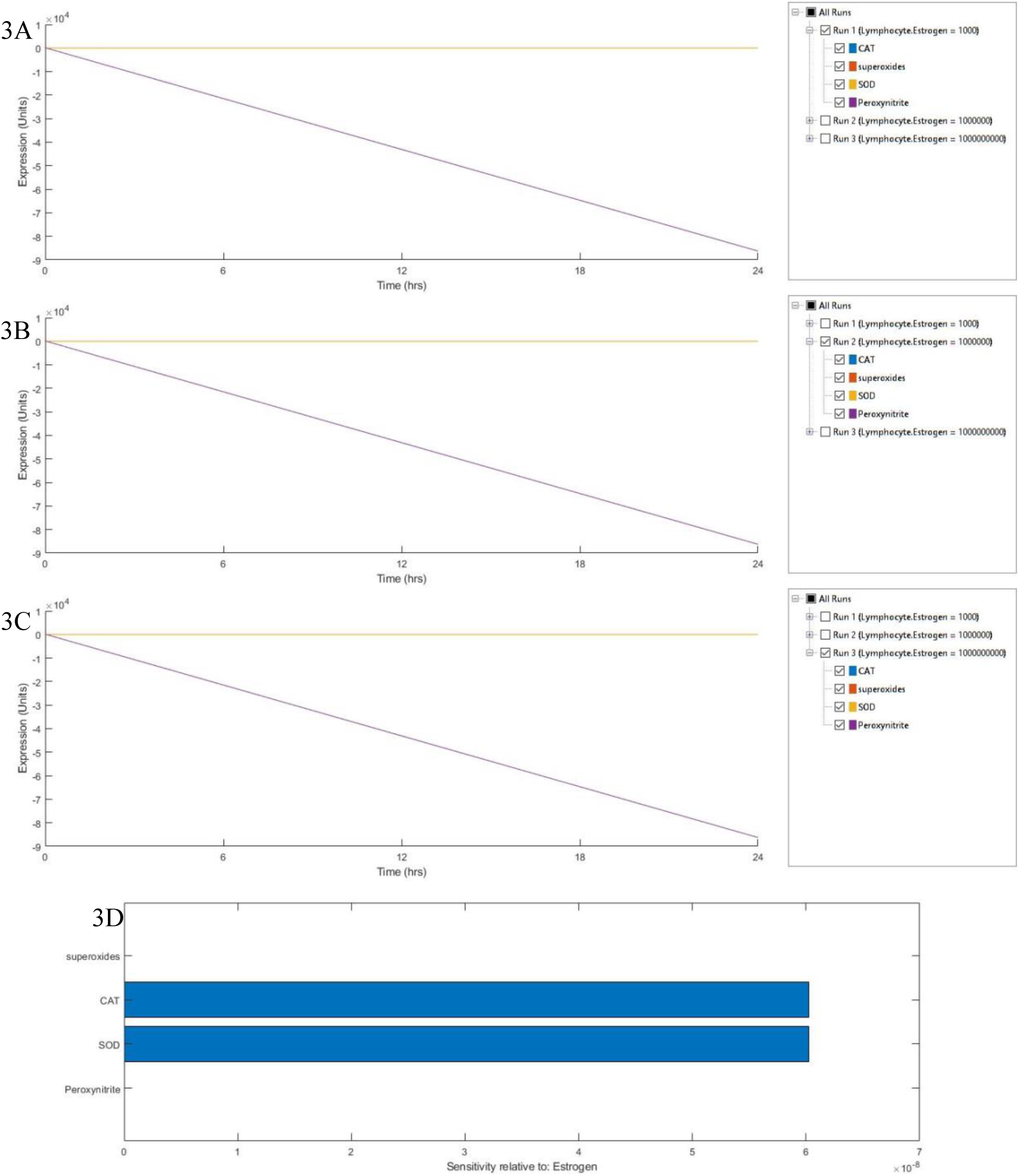
Simulation of Estrogen Signaling: Expression of antioxidant enzymes: Catalase (CAT), Superoxide Dismutase (SOD), and superoxides and peroxynitrites upon stimulation with Estrogen 10^−9^M (A), 10^−6^M (B) and 10^−3^M (C) after 24 hours and their sensitivities relative to Estrogen (D).

**Supplementary Figure 4:**
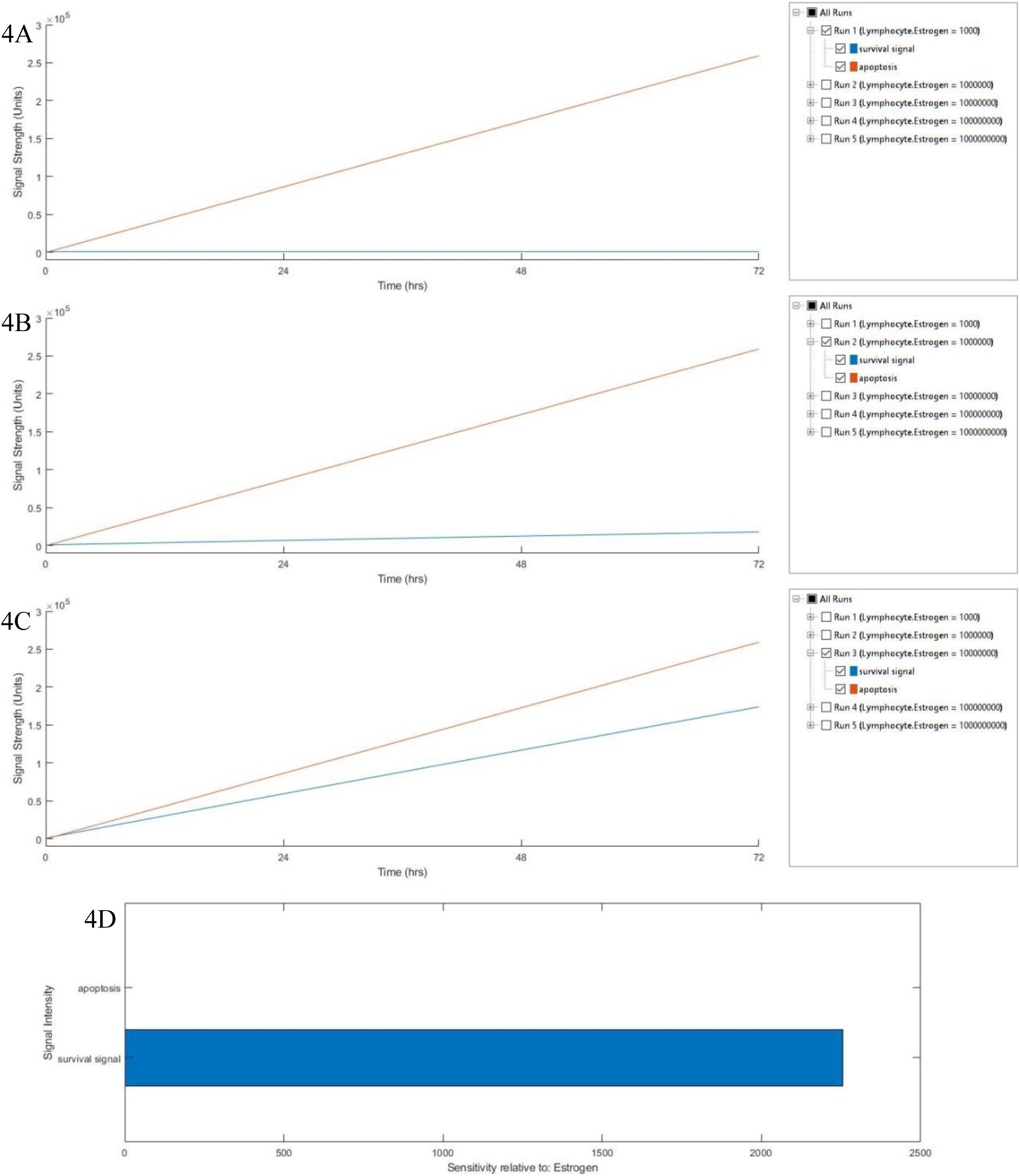
Simulation of Estrogen Signaling: Expression of survival signal and apoptosis upon stimulation with Estrogen 10^−9^M (A), 10^−6^M (B) and 10^−3^M (C) after 72 hours and their sensitivities relative to Estrogen (D).

**Supplementary Figure 5:**
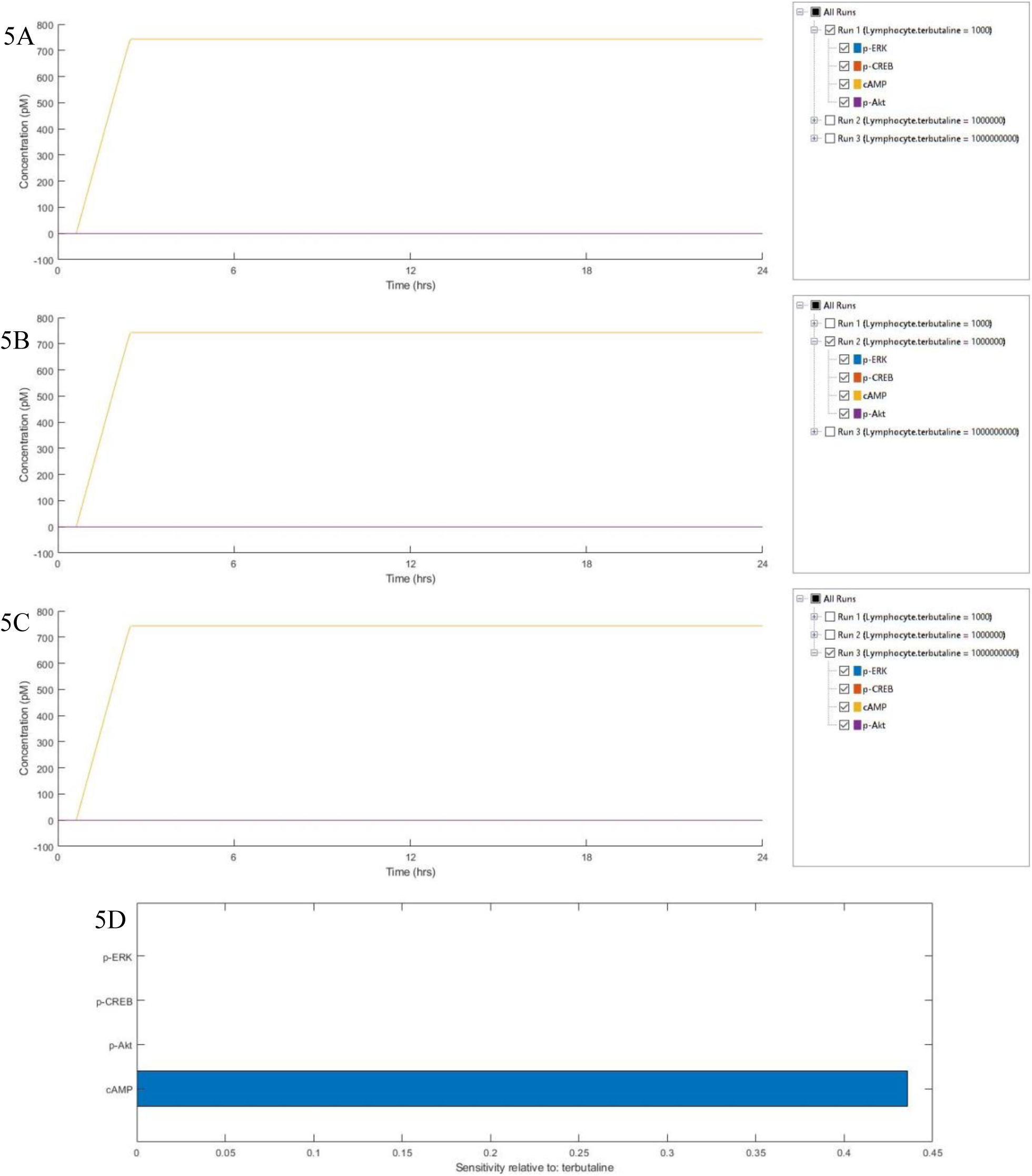
Simulation of Terbutaline Signaling: Expression of molecular markers: p-ERK, p-CREB, cAMP and p-Akt upon stimulation with Terbutaline 10^−9^M (A), 10^−6^M (B) and 10^−3^M (C) after 24 hours and their sensitivities relative to Terbutaline (D).

**Supplementary Figure 6:**
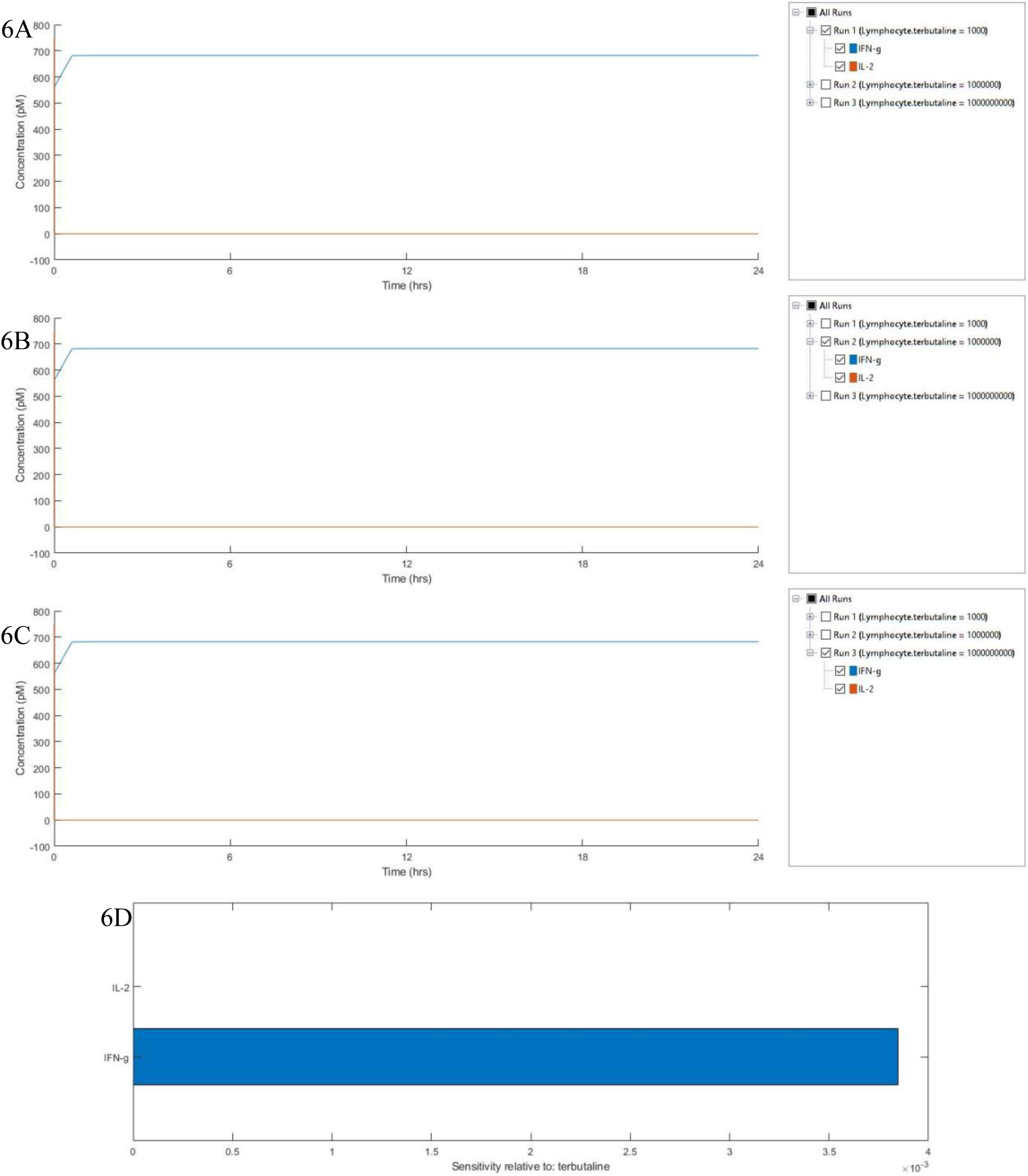
Simulation of Terbutaline Signaling: Expression of cytokines: IFN-γ and IL-2 upon stimulation with Terbutaline 10^−9^M (A), 10^−6^M (B) and 10^−3^M (C) after 24 hours and their sensitivities relative to Terbutaline (D).

**Supplementary Figure 7:**
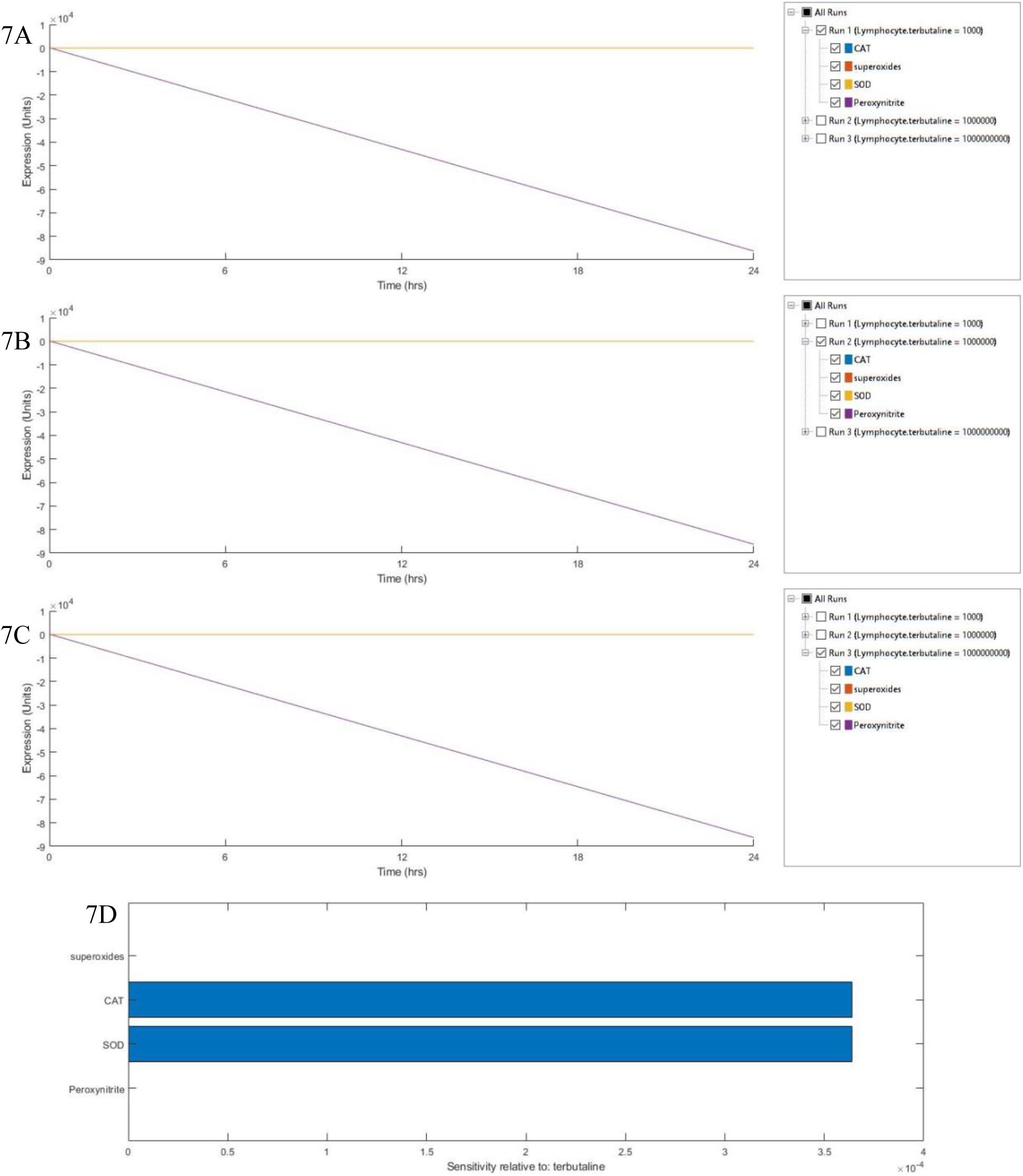
Simulation of Terbutaline Signaling: Expression of antioxidant enzymes: Catalase (CAT), Superoxide Dismutase (SOD), and superoxides and peroxynitrites upon stimulation with Terbutaline 10^−9^M (A), 10^−6^M (B) and 10^−3^M (C) after 24 hours and their sensitivities relative to Terbutaline (D).

**Supplementary Figure 8:**
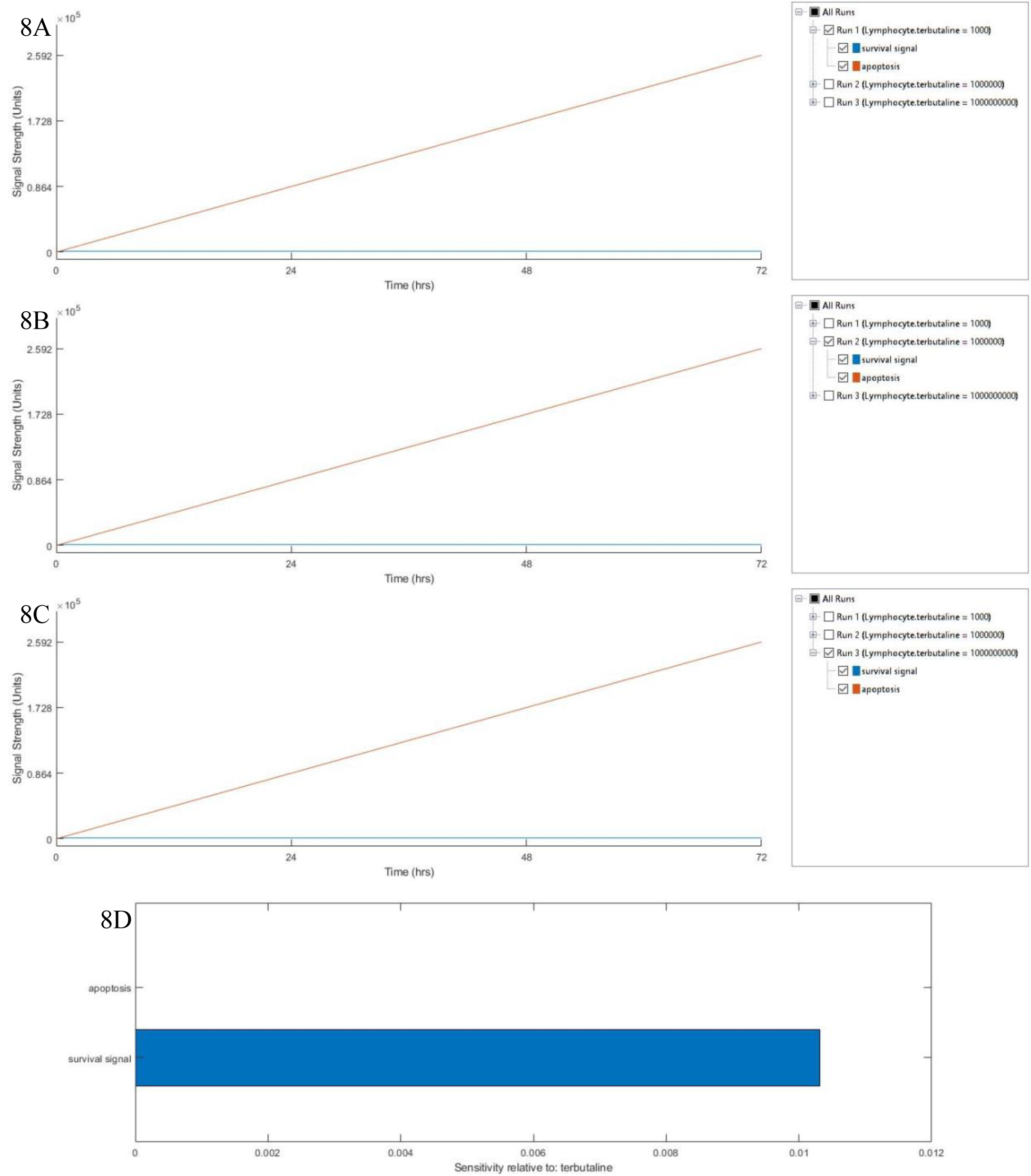
Simulation of Terbutaline Signaling: Expression of survival signal and apoptosis upon stimulation with Terbutaline 10^−9^M (A), 10^−6^M (B) and 10^−3^M (C) after 72 hours and their sensitivities relative to Terbutaline (D).

**Supplementary Figure 9:**
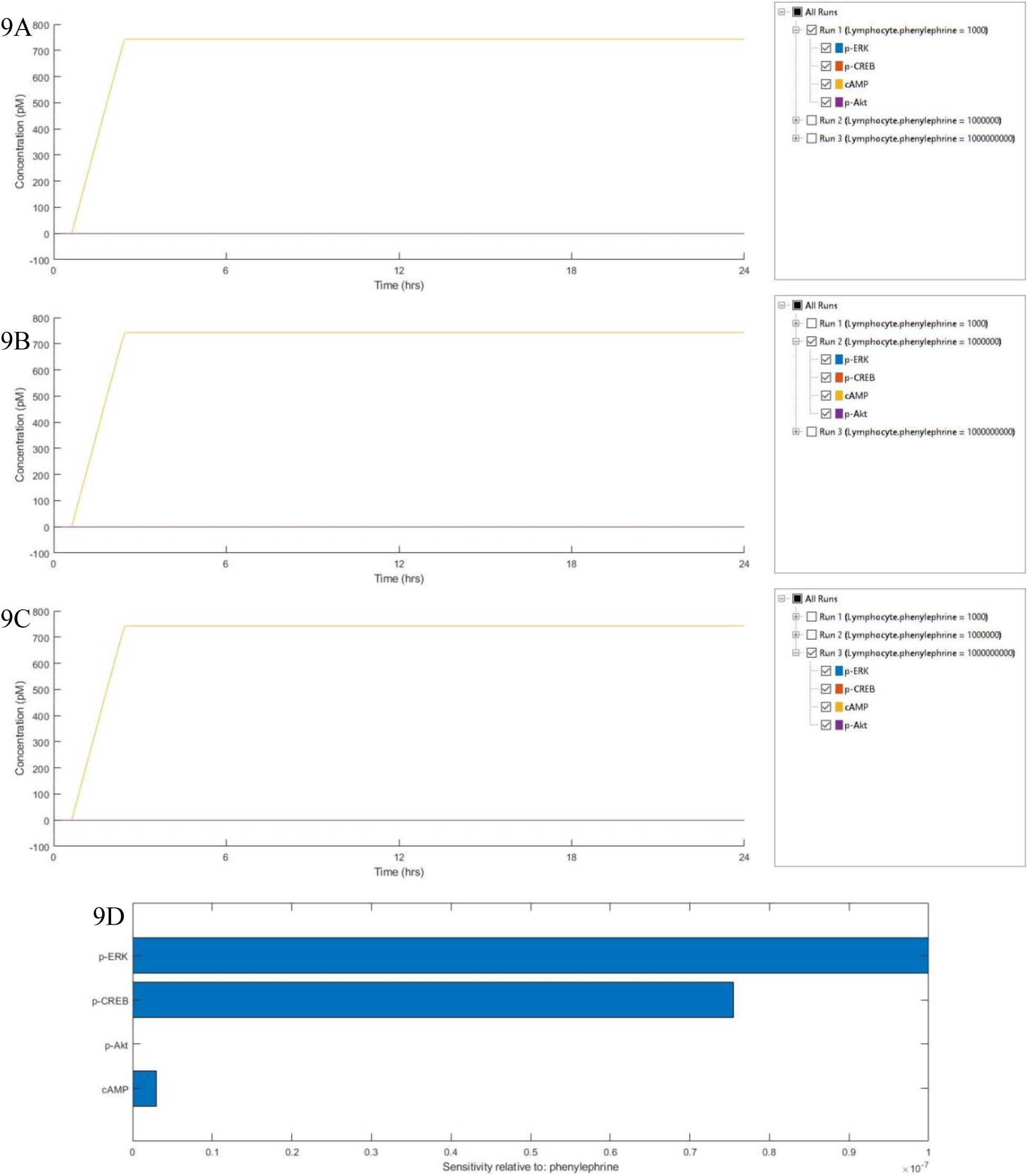
Simulation of Phenylephrine Signaling: Expression of molecular markers: p-ERK, p-CREB, cAMP and p-Akt upon stimulation with Phenylephrine 10^−9^M (A), 10^−6^M (B) and 10^−3^M (C) after 24 hours and their sensitivities relative to Phenylephrine (D).

**Supplementary Figure 10:**
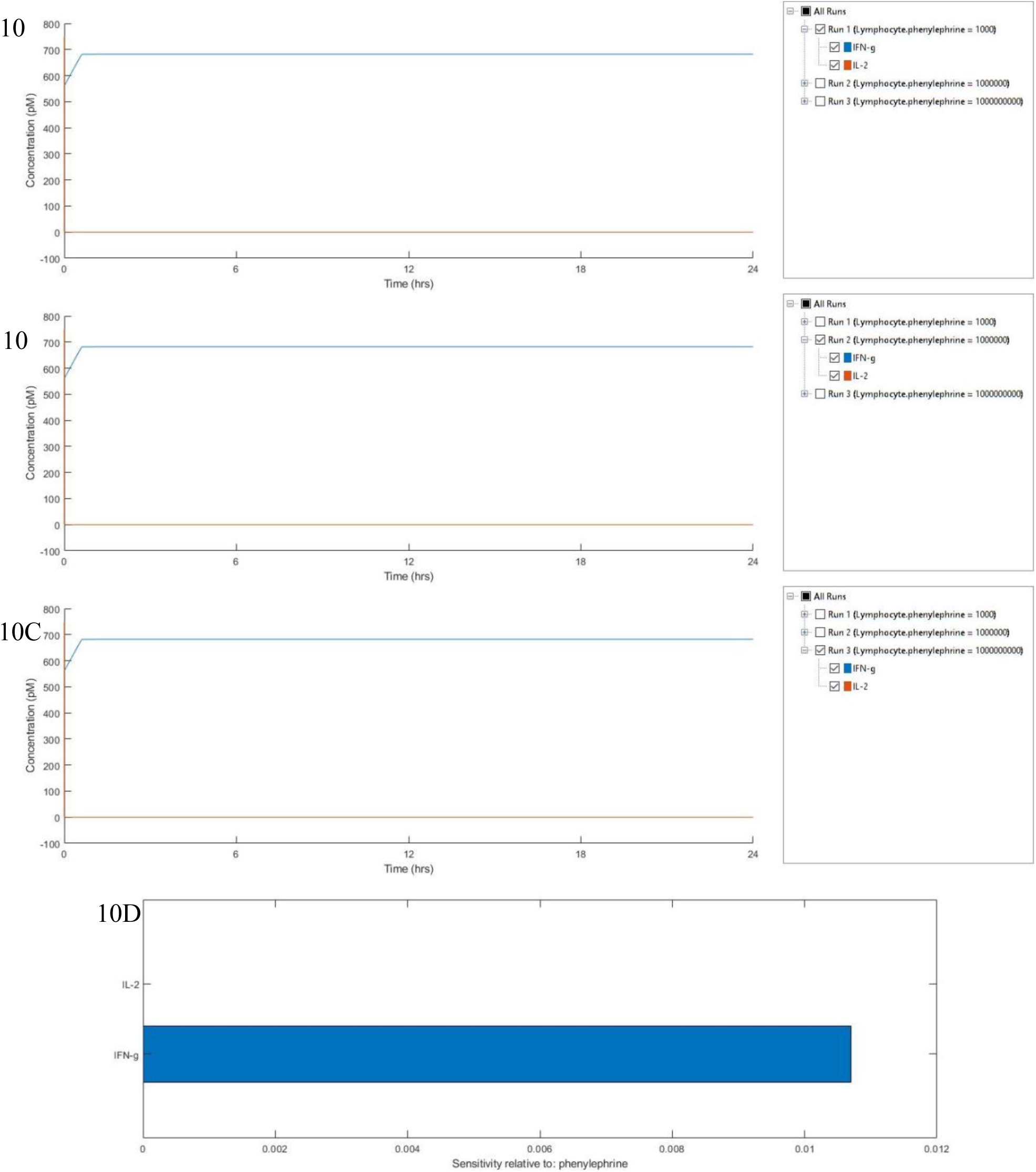
Simulation of Phenylephrine Signaling: Expression of cytokines: IFN-γ and IL-2 upon stimulation with Phenylephrine 10^−9^M (A), 10^−6^M (B) and 10^−3^M (C) after 24 hours and their sensitivities relative to Phenylephrine (D).

**Supplementary Figure 11:**
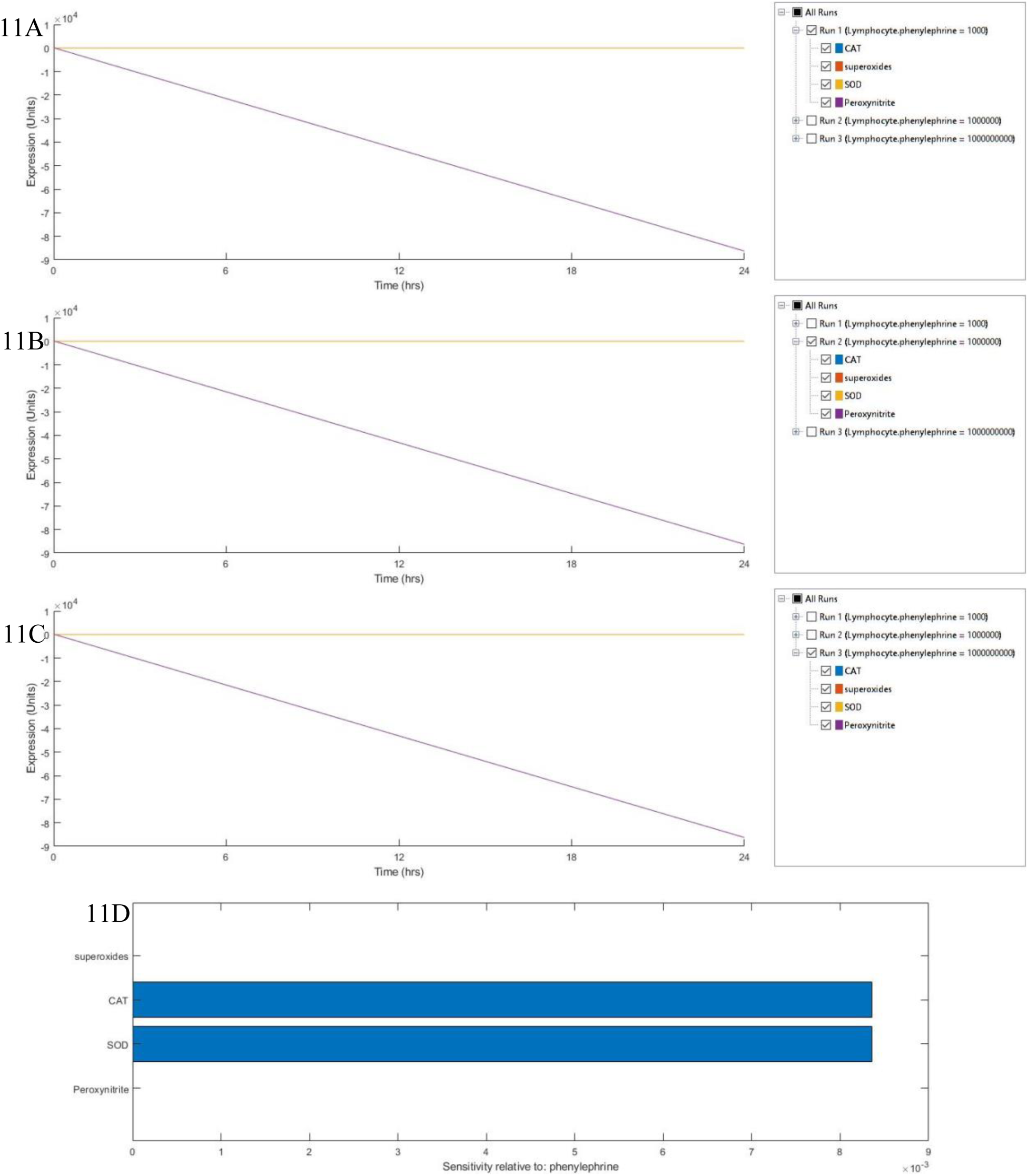
Simulation of Phenylephrine Signaling: Expression of antioxidant enzymes: Catalase (CAT), Superoxide Dismutase (SOD), and superoxides and peroxynitrites upon stimulation with Phenylephrine 10^−9^M (A), 10^−6^M (B) and 10^−3^M (C) after 24 hours and their sensitivities relative to Phenylephrine (D).

**Supplementary Figure 12:**
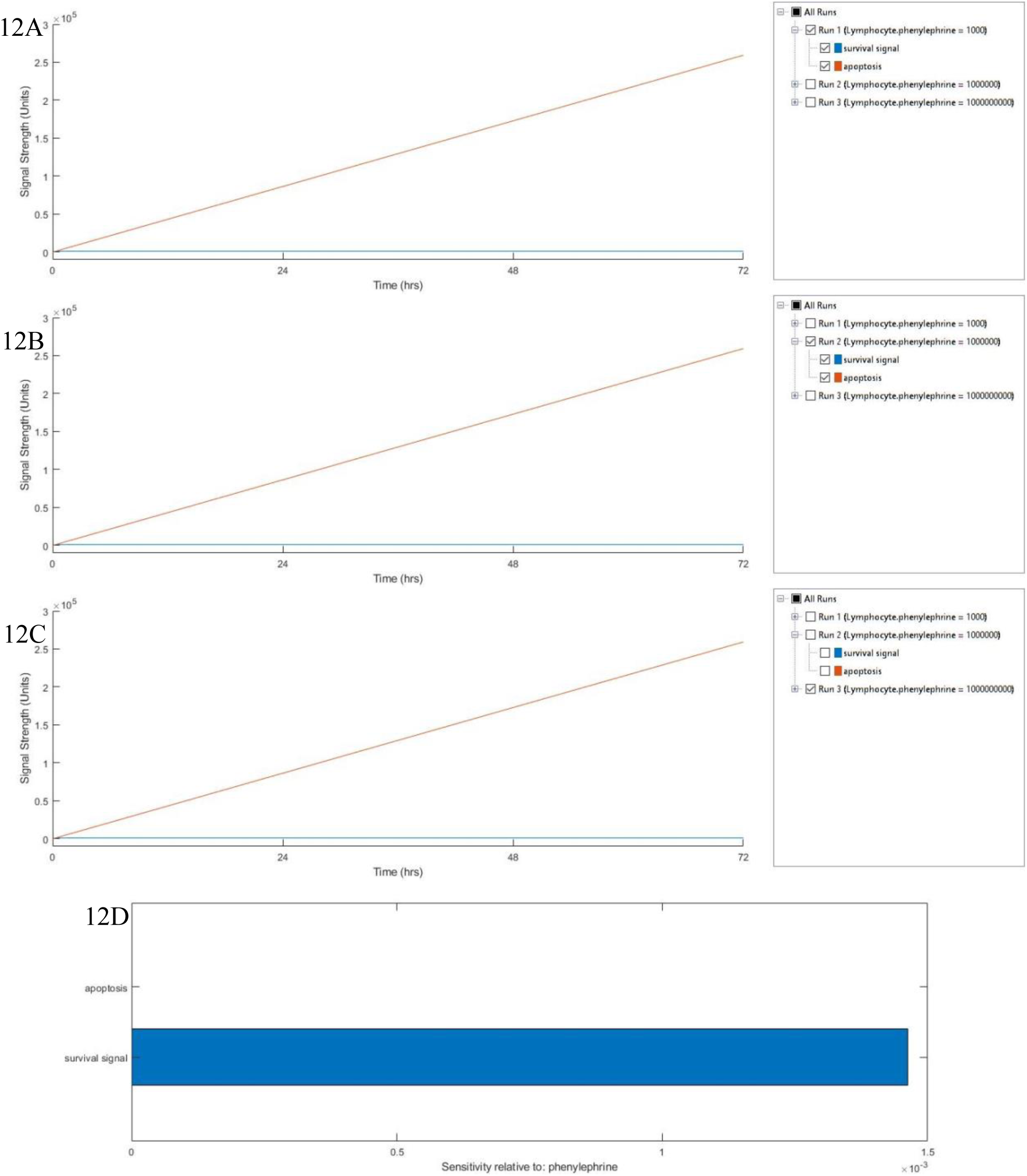
Simulation of Phenylephrine Signaling: Expression of survival signal and apoptosis upon stimulation with Phenylephrine 10^−9^M (A), 10^−6^M (B) and 10^−3^M (C) after 72 hours and their sensitivities relative to Phenylephrine (D).

**Supplementary Figure 13:**
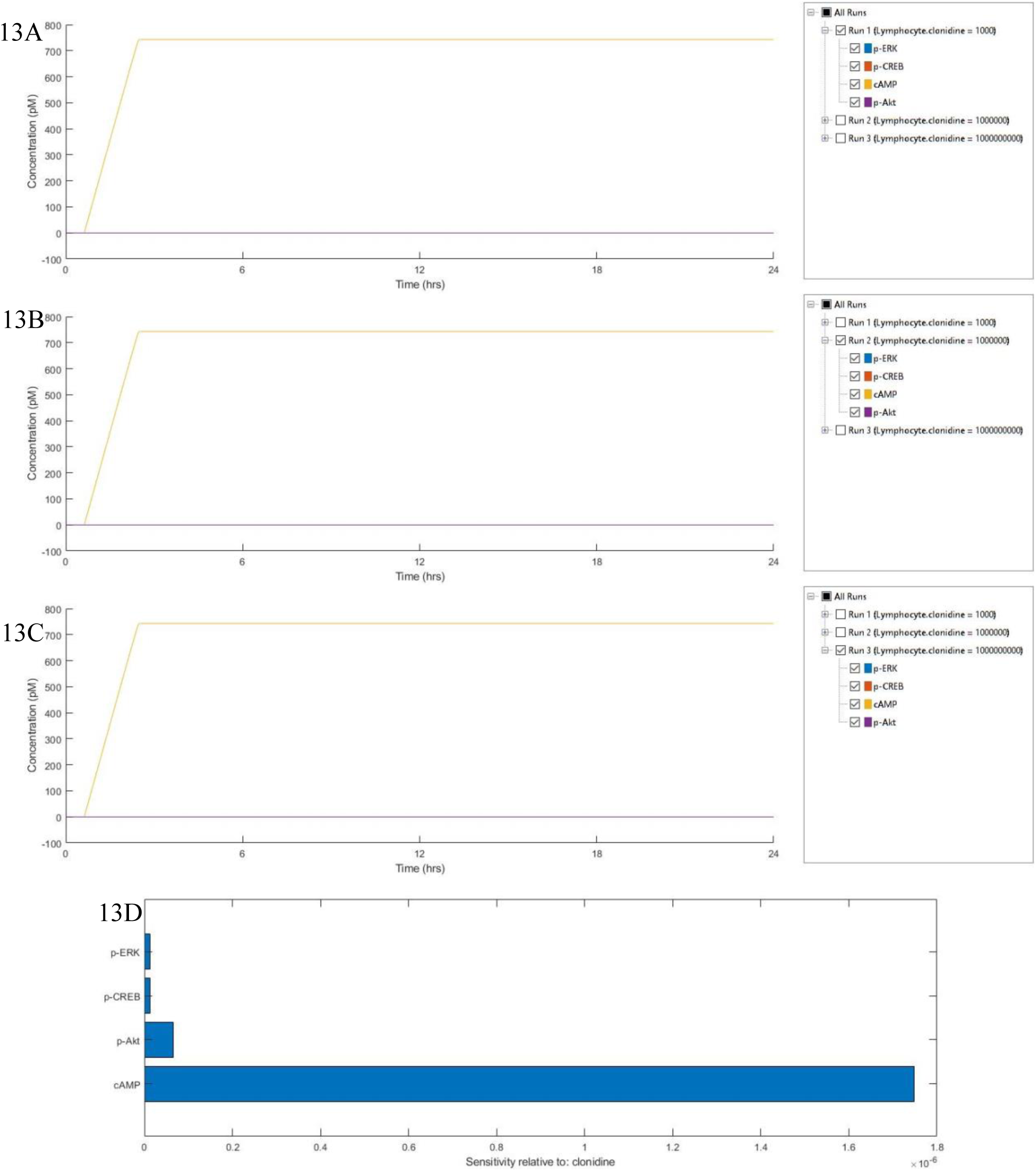
Simulation of Clonidine Signaling: Expression of molecular markers: p-ERK, p-CREB, cAMP and p-Akt upon stimulation with Clonidine 10^−9^M (A), 10^−6^M (B) and 10^−3^M (C) after 24 hours and their sensitivities relative to Clonidine (D).

**Supplementary Figure 14:**
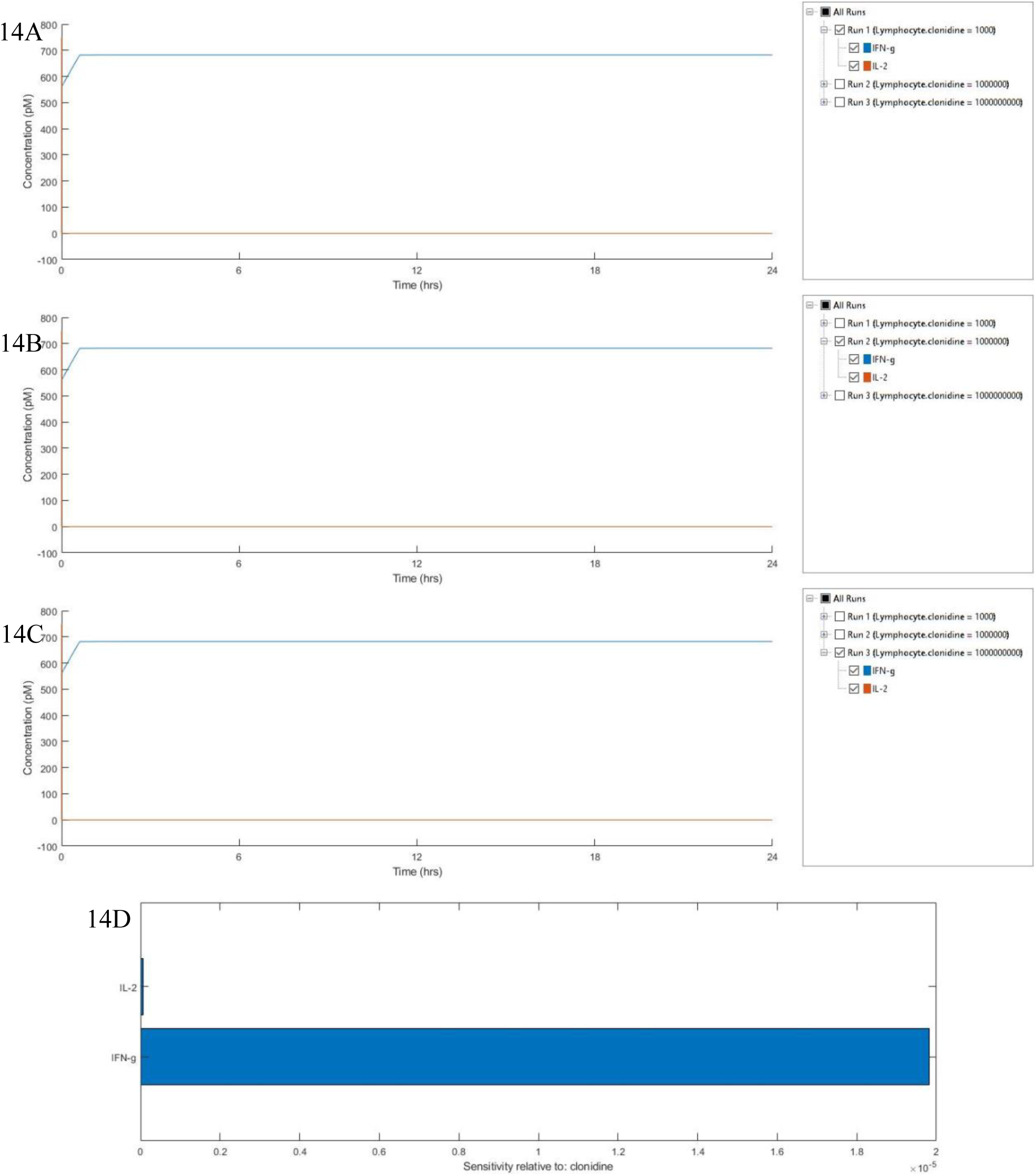
Simulation of Clonidine Signaling: Expression of cytokines: IFN-γ and IL-2 upon stimulation with Clonidine 10^−9^M (A), 10^−6^M (B) and 10^−3^M (C) after 24 hours and their sensitivities relative to Clonidine (D).

**Supplementary Figure 15:**
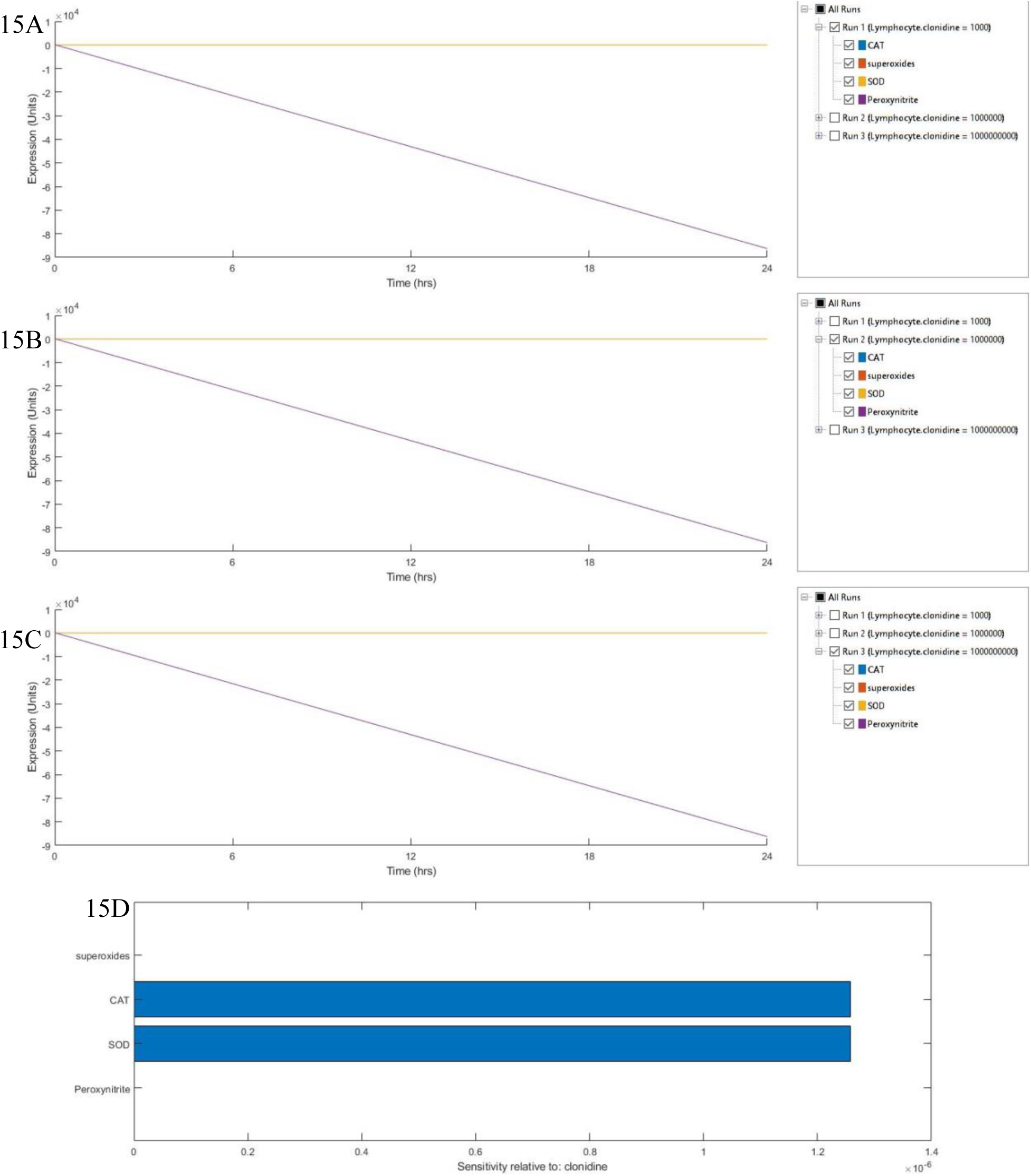
Simulation of Clonidine Signaling: Expression of antioxidant enzymes: Catalase (CAT), Superoxide Dismutase (SOD), and superoxides and peroxynitrites upon stimulation with Clonidine 10^−9^M (A), 10^−6^M (B) and 10^−3^M (C) after 24 hours and their sensitivities relative to Clonidine (D).

**Supplementary Figure 16:**
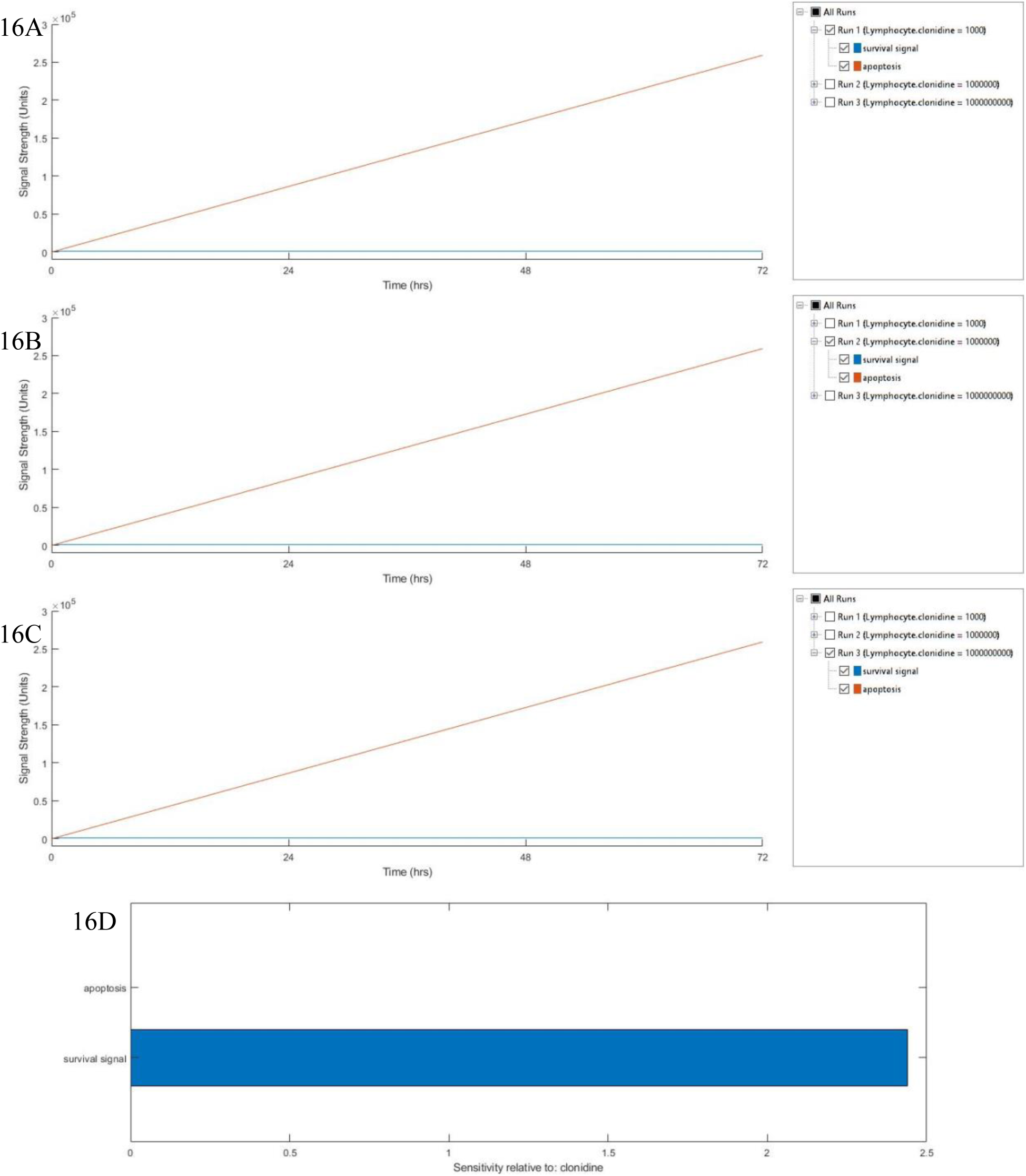
Simulation of Clonidine Signaling: Expression of survival signal and apoptosis upon stimulation with Clonidine 10^−9^M (A), 10^−6^M (B) and 10^−3^M (C) after 72 hours and their sensitivities relative to Clonidine (D).

**Supplementary Figure 17:**
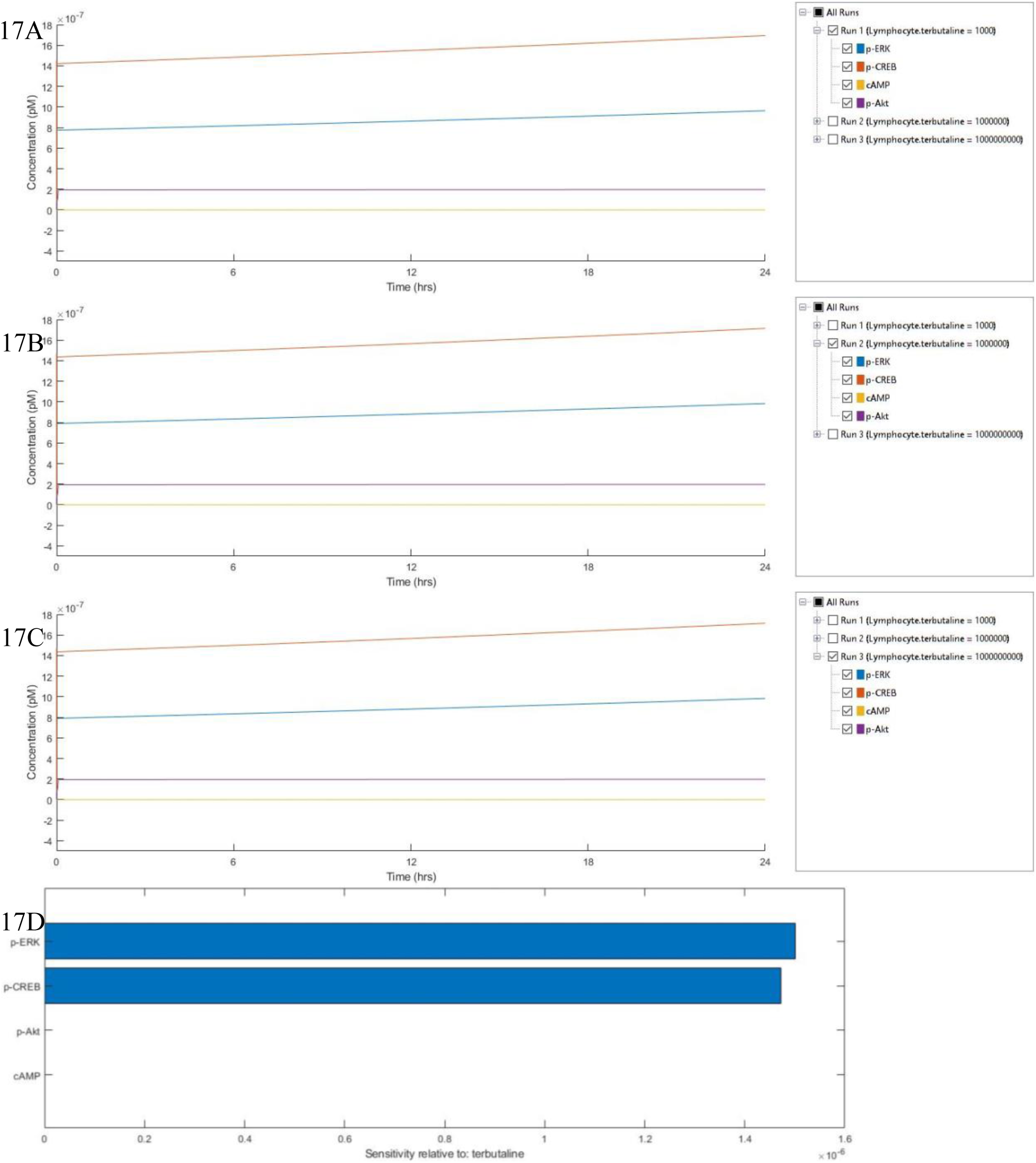
Simulation of Terbutaline+Estrogen Signaling: Expression of molecular markers: p-ERK, p-CREB, cAMP and p-Akt upon stimulation with Terbutaline 10^−9^M (A), 10^−6^M (B) and 10^−3^M (C) +Estrogen 10^−6^M after 24 hours and their sensitivities relative to Terbutaline+Estrogen (D).

**Supplementary Figure 18:**
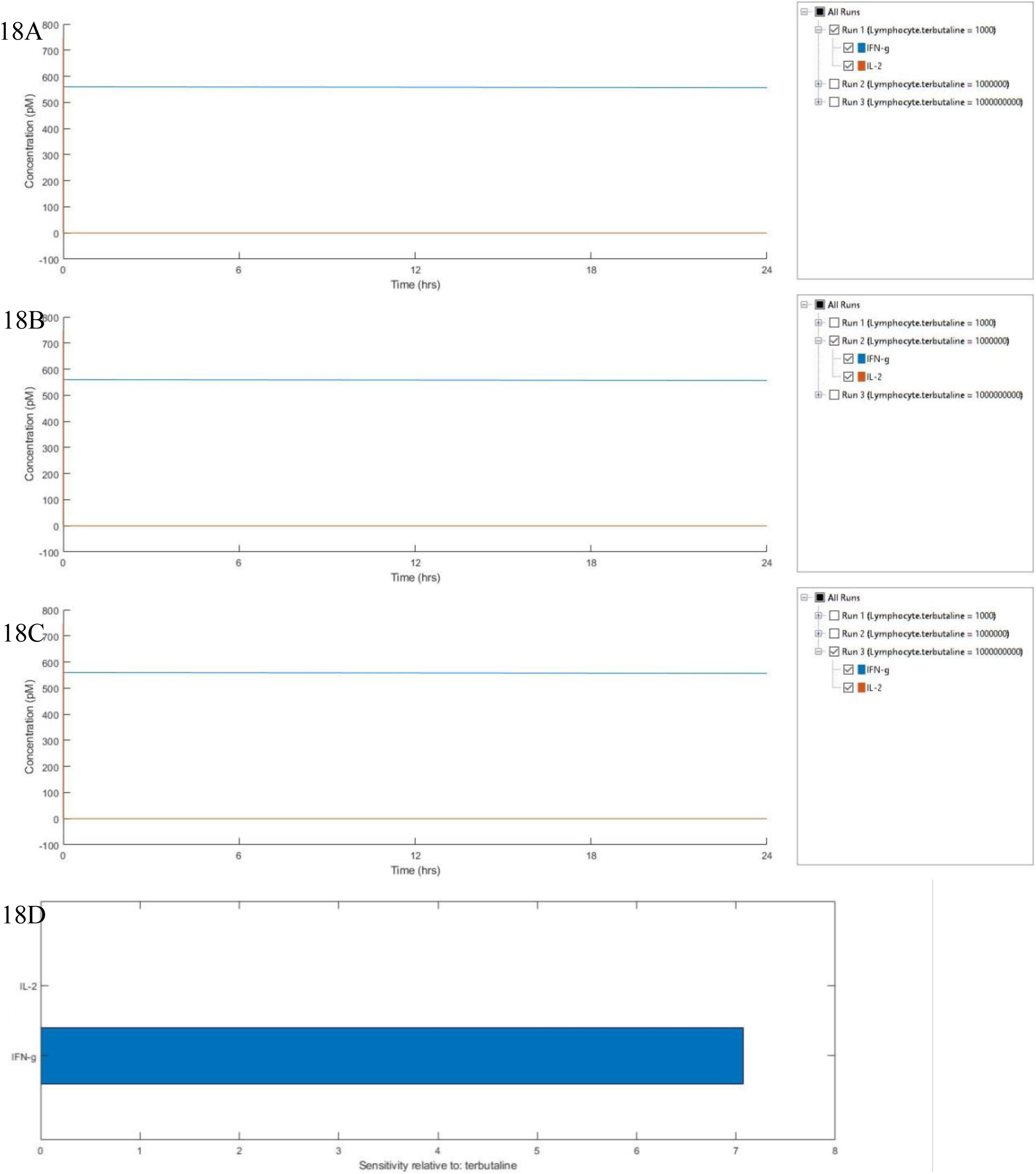
Simulation of Terbutaline+Estrogen Signaling: Expression of cytokines: IFN-γ and IL-2 upon stimulation with Terbutaline 10^−9^M (A), 10^−6^M (B) and 10^−3^M (C) +Estrogen 10^−6^M after 24 hours and their sensitivities relative to Terbutaline+Estrogen (D).

**Supplementary Figure 19:**
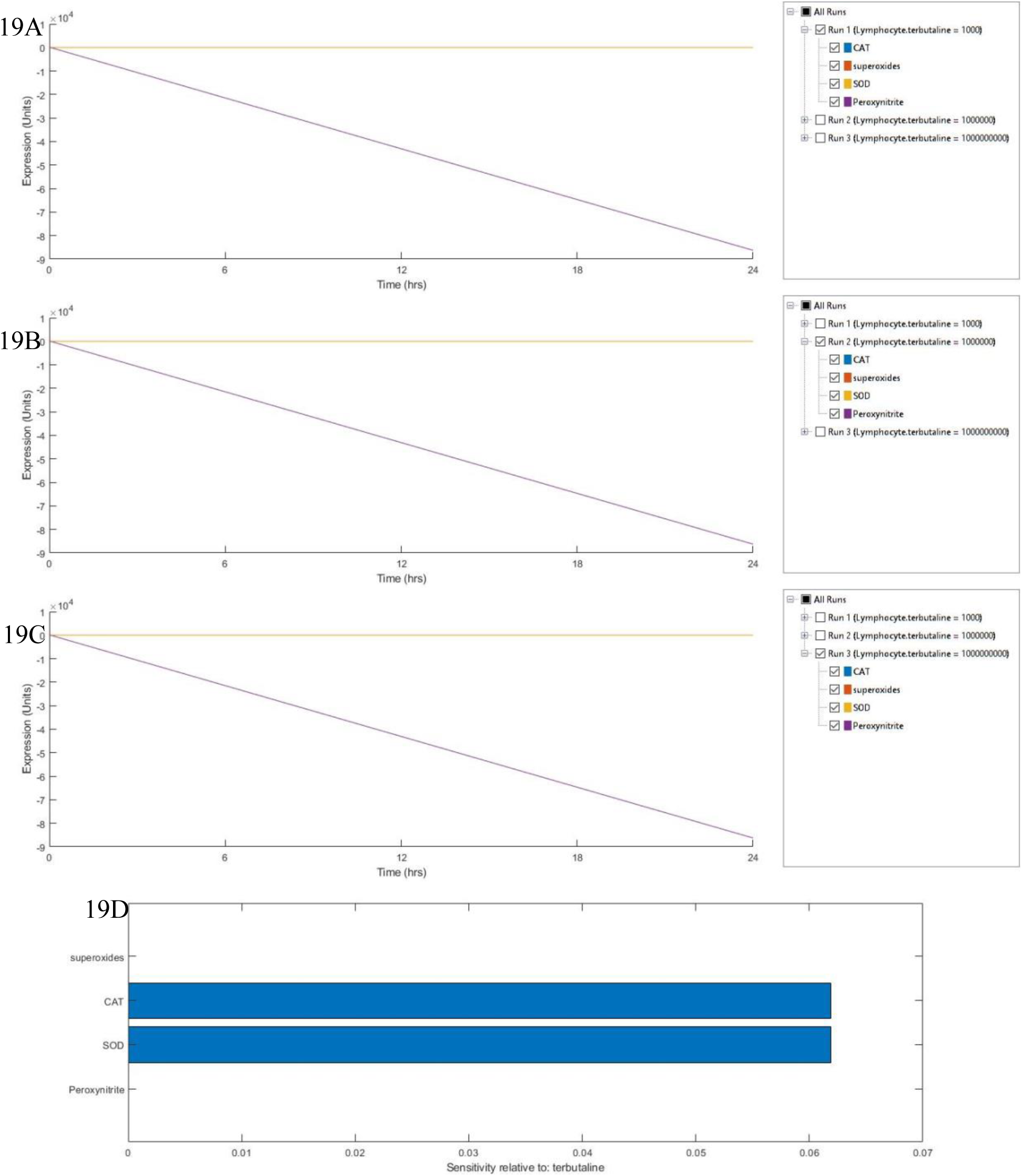
Simulation of Terbutaline+Estrogen Signaling: Expression of antioxidant enzymes: Catalase (CAT), Superoxide Dismutase (SOD), and superoxides and peroxynitrites upon stimulation with Terbutaline 10^−9^M (A), 10^−6^M (B) and 10^−3^M (C) +Estrogen 10^−6^M after 24 hours and their sensitivities relative to Terbutaline+Estrogen (D).

**Supplementary Figure 20:**
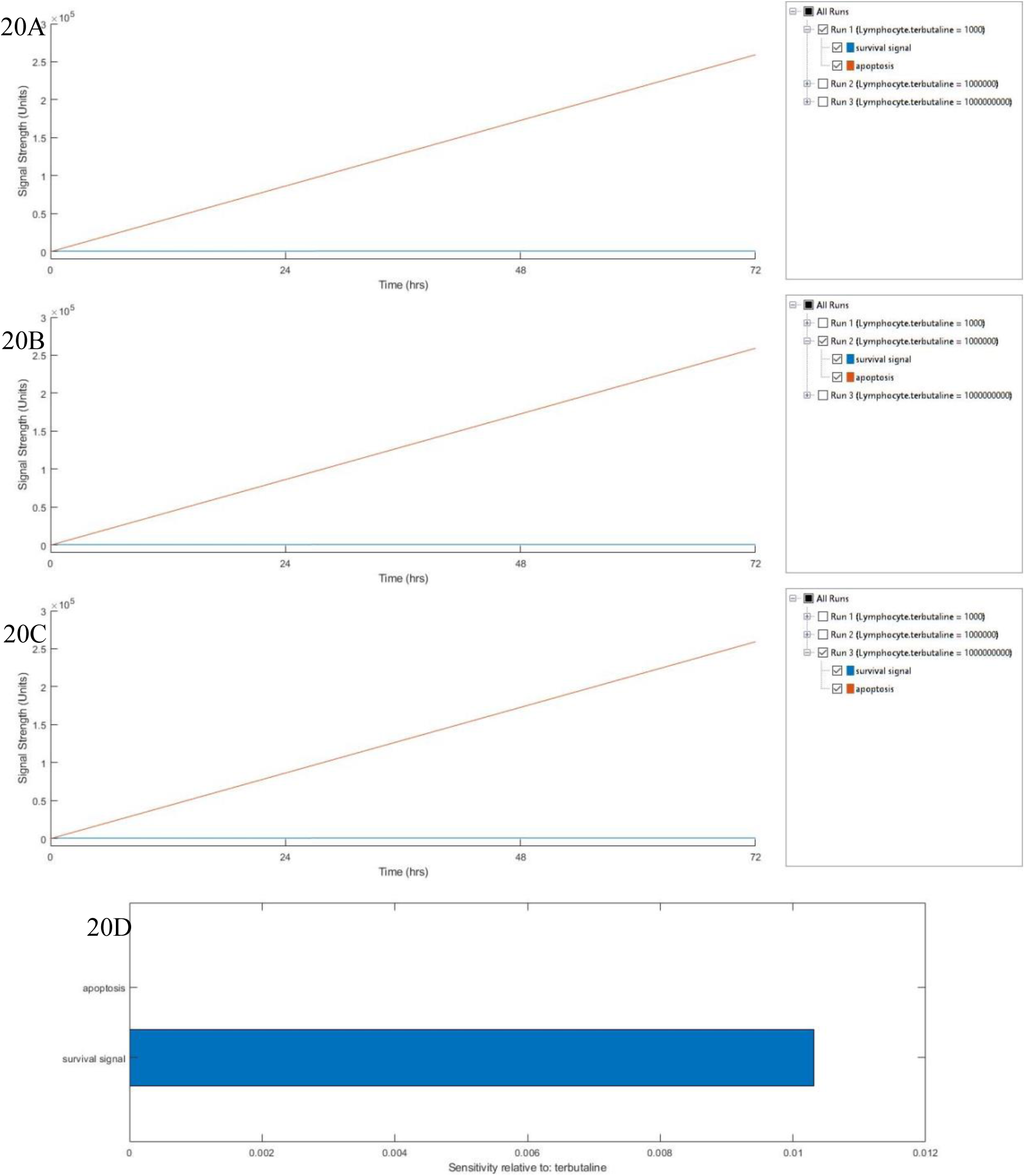
Simulation of Terbutaline+Estrogen Signaling: Expression of survival signal and apoptosis upon stimulation with Terbutaline 10^−9^M (A), 10^−6^M (B) and 10^−3^M (C) +Estrogen 10^−6^M after 72 hours and their sensitivities relative to Terbutaline+Estrogen (D).

**Supplementary Figure 21:**
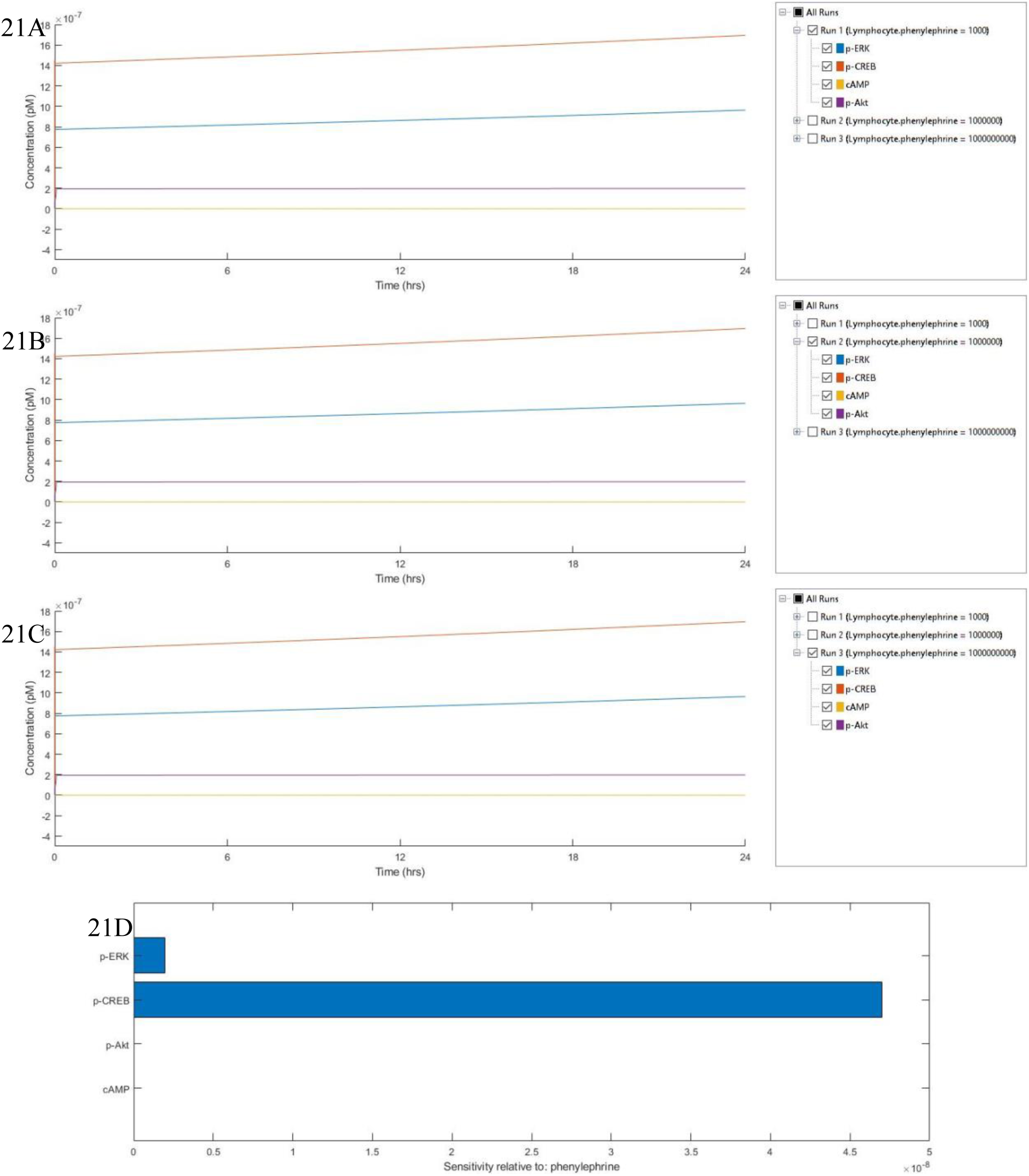
Simulation of Phenylephrine+Estrogen Signaling: Expression of molecular markers: p-ERK, p-CREB, cAMP and p-Akt upon stimulation with Phenylephrine 10^−9^M (A), 10^−6^M (B) and 10^−3^M (C) +Estrogen 10^−6^M after 24 hours and their sensitivities relative to Phenylephrine+Estrogen (D).

**Supplementary Figure 22:**
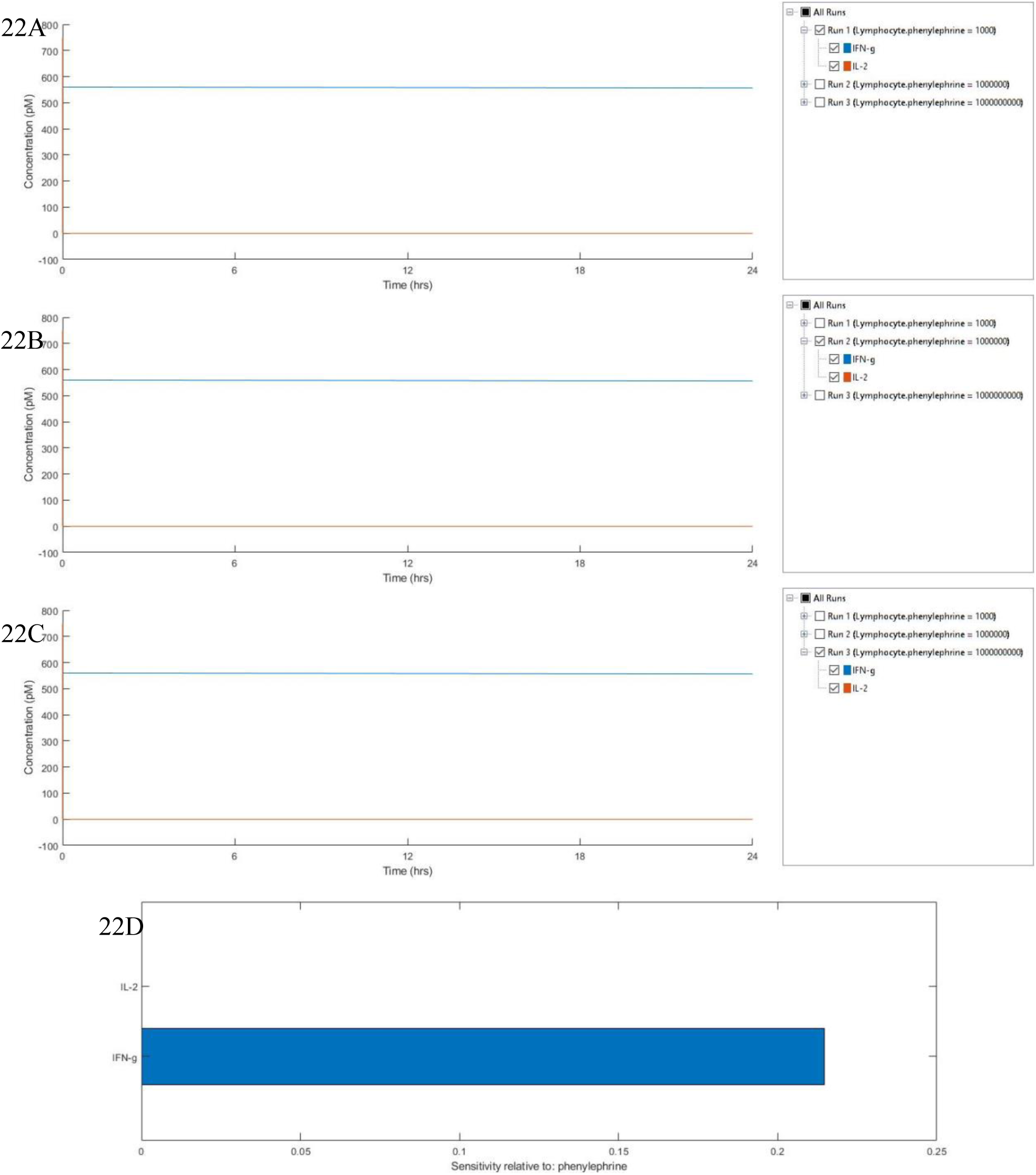
Simulation of Phenylephrine+Estrogen Signaling: Expression of cytokines: IFN-γ and IL-2 upon stimulation with Phenylephrine 10^−9^M (A), 10^−6^M (B) and 10^−3^M (C) +Estrogen 10^−6^M after 24 hours and their sensitivities relative to Phenylephrine+Estrogen (D).

**Supplementary Figure 23:**
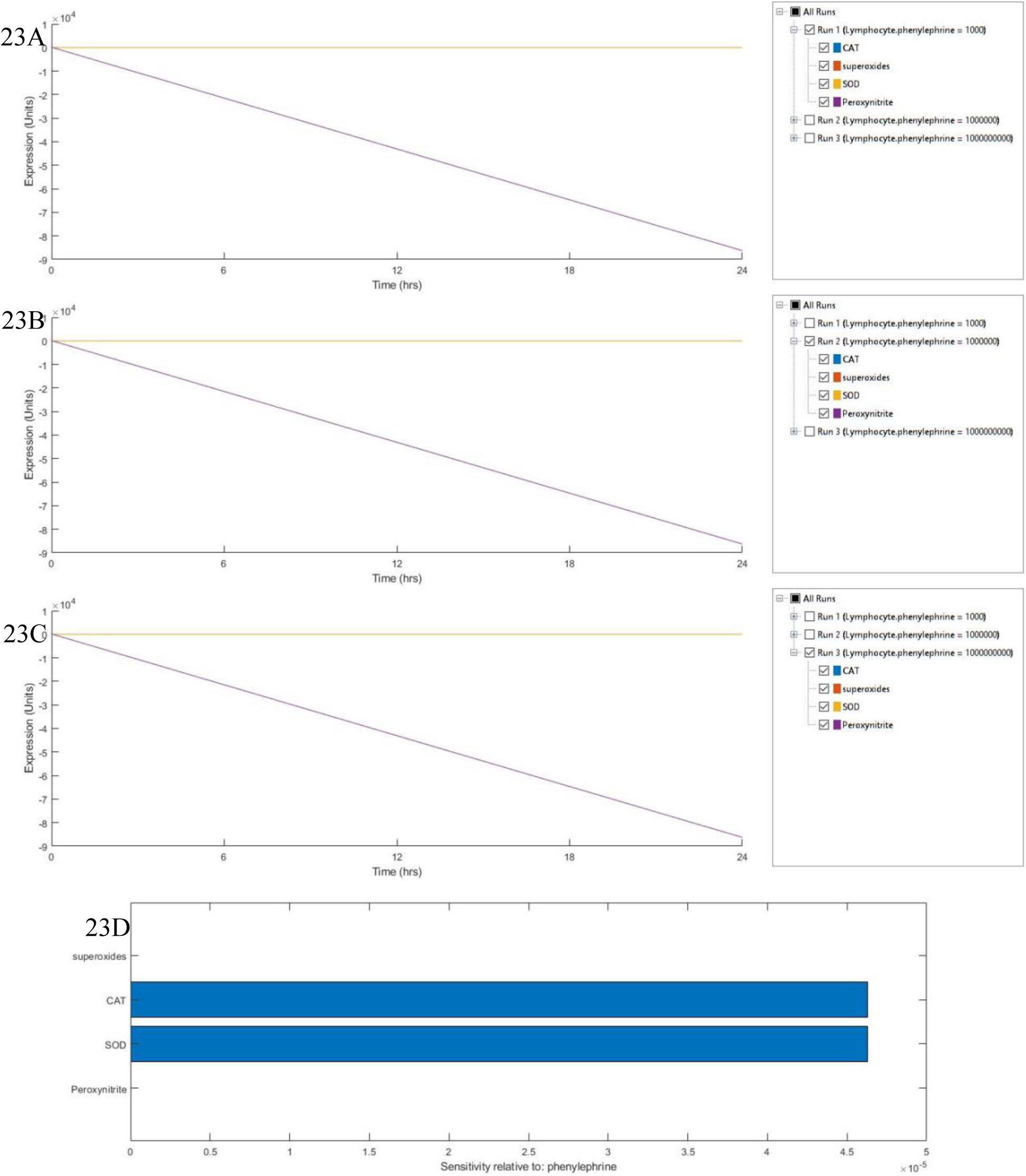
Simulation of Phenylephrine+Estrogen Signaling: Expression of antioxidant enzymes: Catalase (CAT), Superoxide Dismutase (SOD), and superoxides and peroxynitrites upon stimulation with Phenylephrine 10^−9^M (A), 10^−6^M (B) and 10^−3^M (C) +Estrogen 10^−6^M after 24 hours and their sensitivities relative to Phenylephrine+Estrogen (D).

**Supplementary Figure 24:**
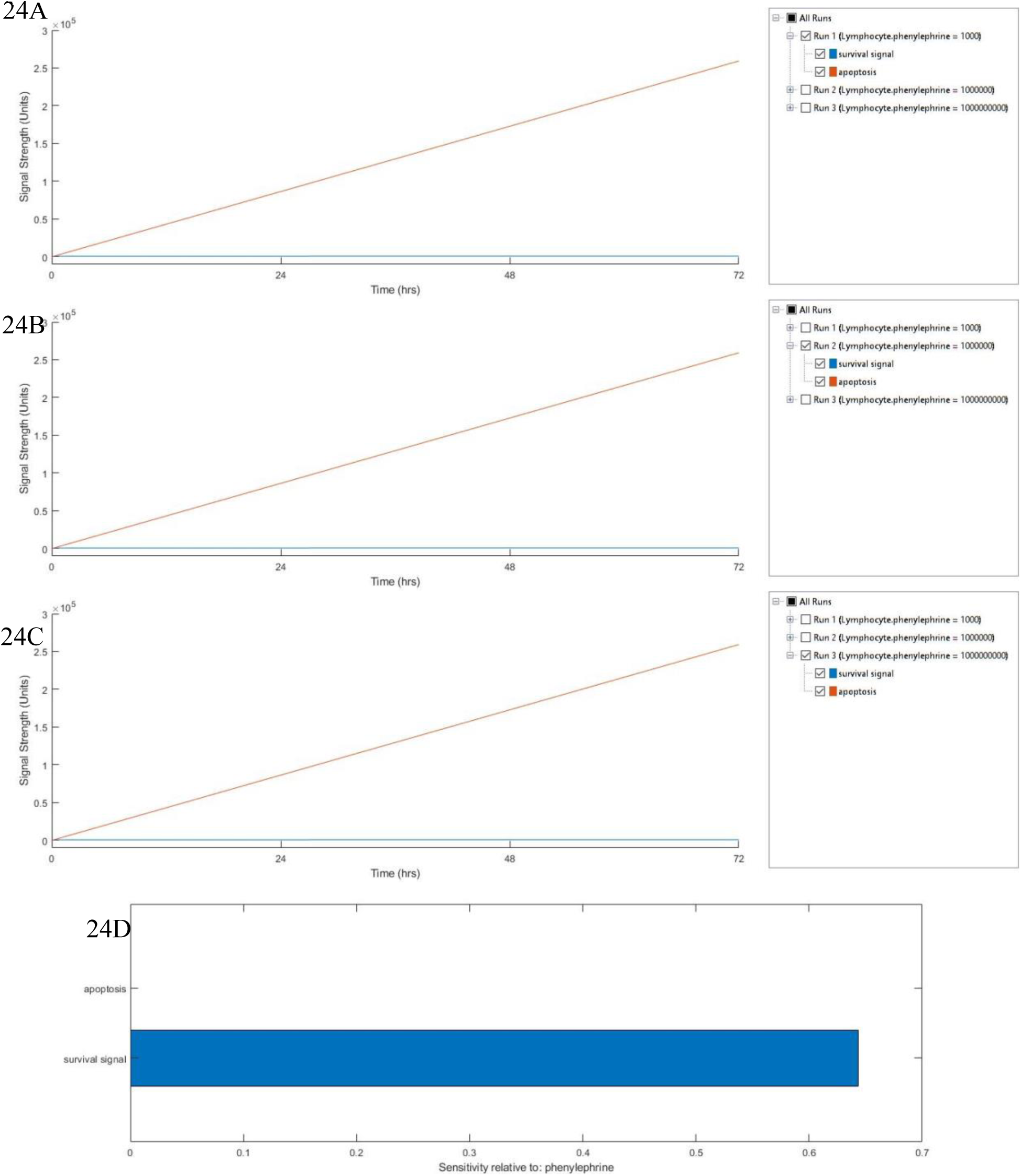
Simulation of Phenylephrine+Estrogen Signaling: Expression of survival signal and apoptosis upon stimulation with Phenylephrine 10^−9^M (A), 10^−6^M (B) and 10^−3^M (C) +Estrogen 10^−6^M after 72 hours and their sensitivities relative to Phenylephrine+Estrogen (D).

**Supplementary Figure 25:**
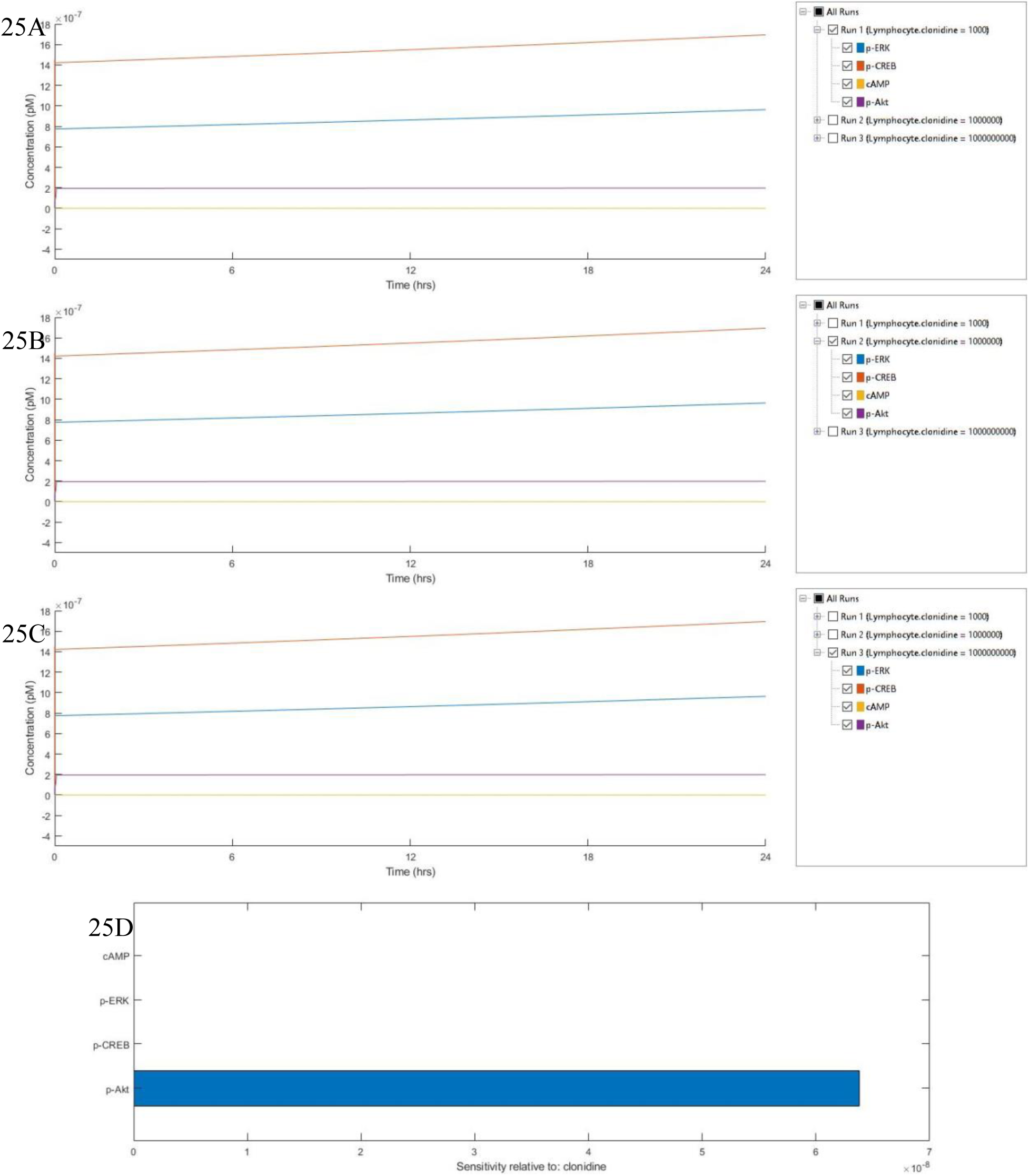
Simulation of Clonidine+Estrogen Signaling: Expression of molecular markers: p-ERK, p-CREB, cAMP and p-Akt upon stimulation with Clonidine 10^−9^M (A), 10^−6^M (B) and 10^−3^M (C) +Estrogen 10^−6^M after 24 hours and their sensitivities relative to Clonidine+Estrogen (D).

**Supplementary Figure 26:**
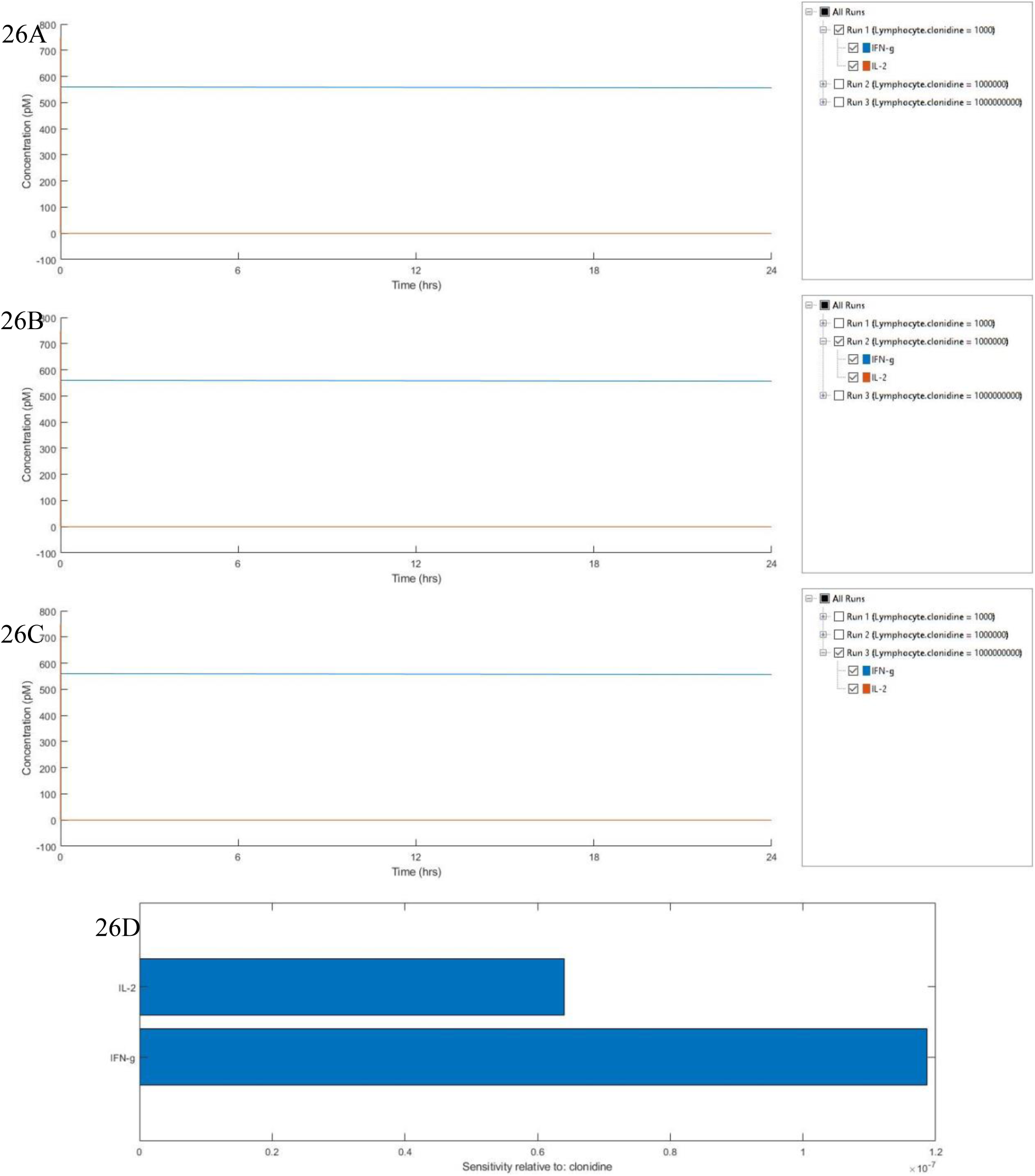
Simulation of Clonidine+Estrogen Signaling: Expression of cytokines: IFN-γ and IL-2 upon stimulation with Clonidine 10^−9^M (A), 10^−6^M (B) and 10^−3^M (C) +Estrogen 10^−6^M after 24 hours and their sensitivities relative to Clonidine+Estrogen (D).

**Supplementary Figure 27:**
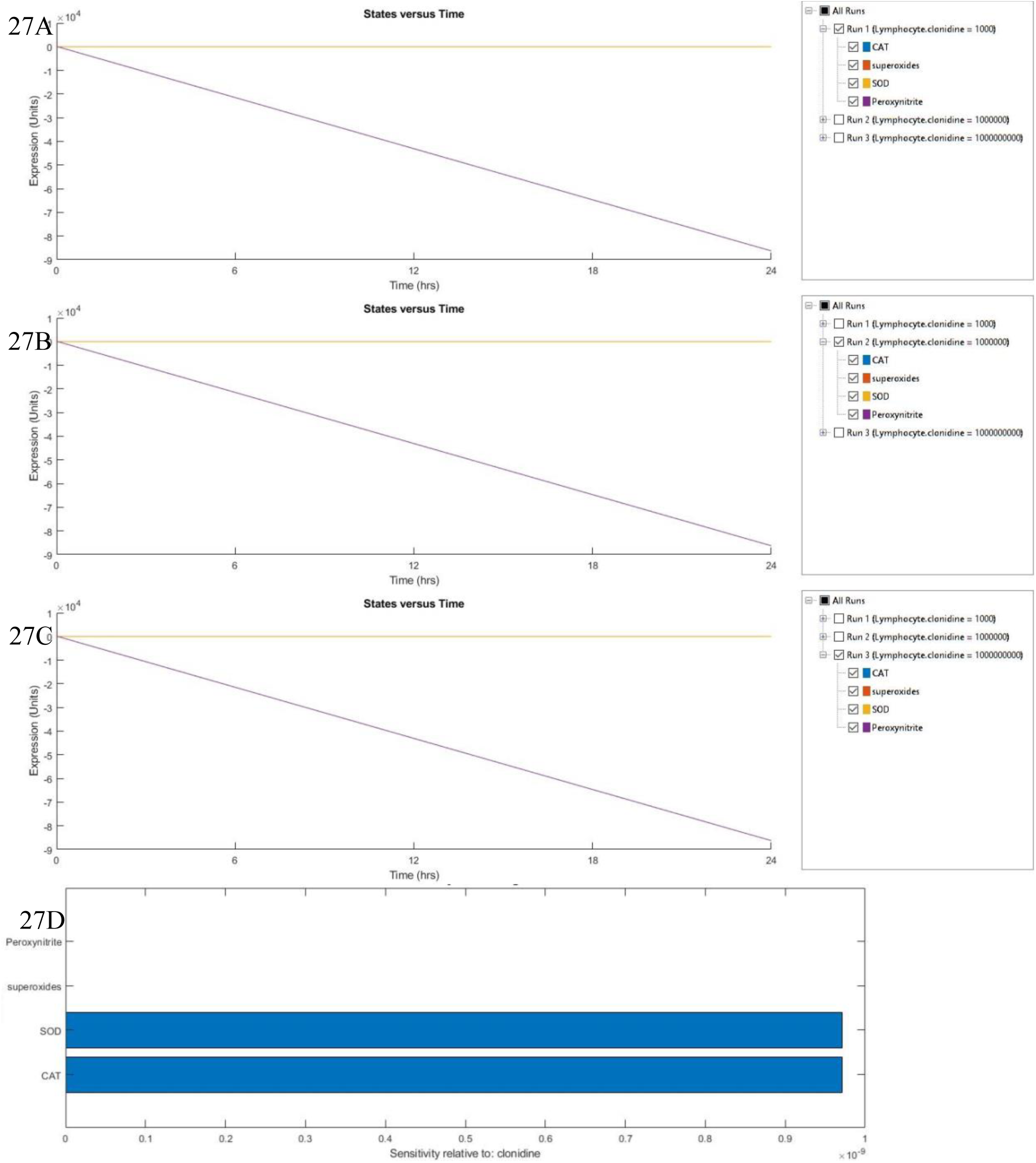
Simulation of Clonidine+Estrogen Signaling: Expression of antioxidant enzymes: Catalase (CAT), Superoxide Dismutase (SOD), and superoxides and peroxynitrites upon stimulation with Clonidine 10^−9^M (A), 10^−6^M (B) and 10^−3^M (C) +Estrogen 10^−6^M after 24 hours and their sensitivities relative to Clonidine+Estrogen (D).

**Supplementary Figure 28:**
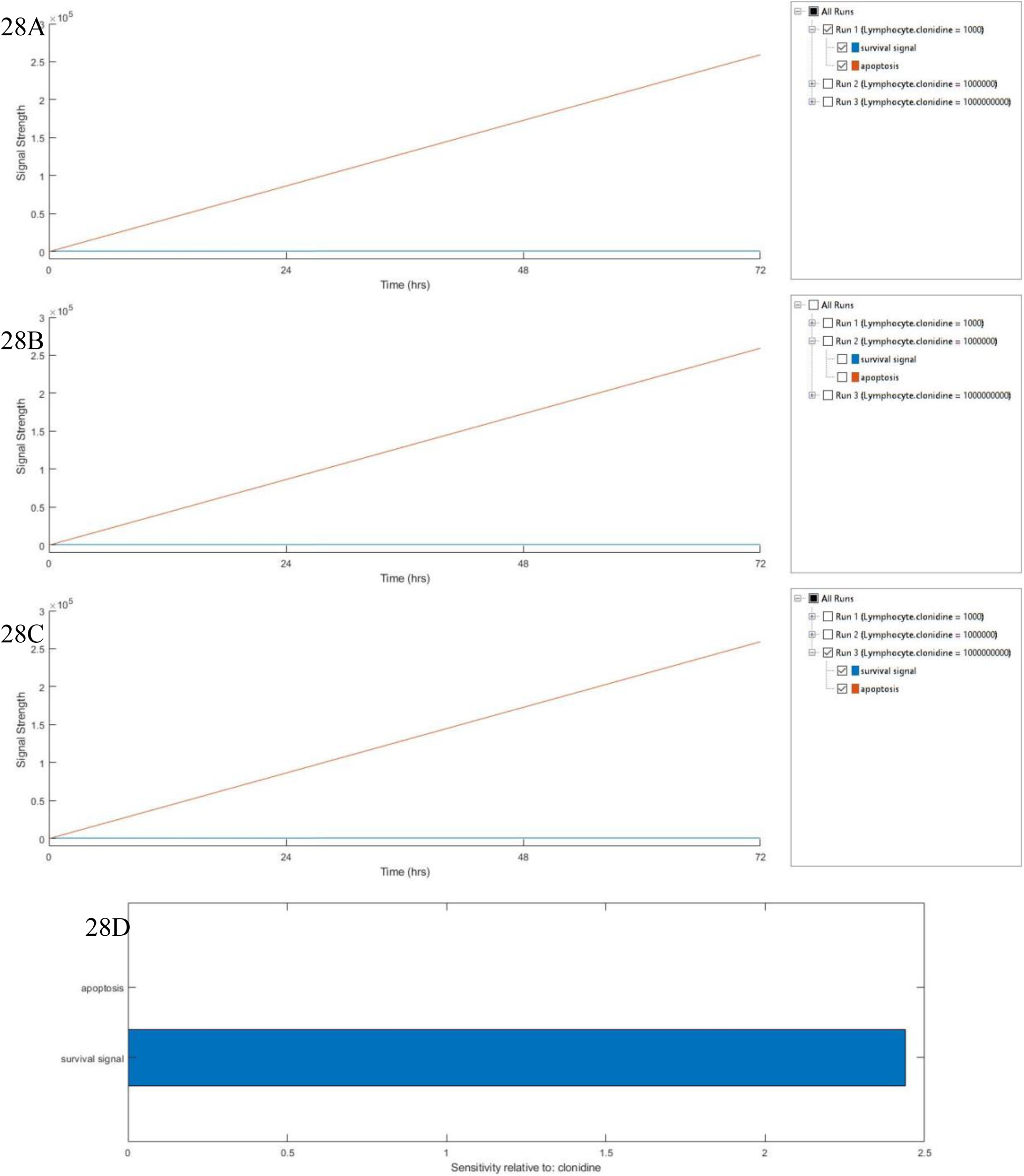
Simulation of Clonidine+Estrogen Signaling: Expression of survival signal and apoptosis upon stimulation with Clonidine 10^−9^M (A), 10^−6^M (B) and 10^−3^M (C) +Estrogen after 72 hours and their sensitivities relative to Clonidine+Estrogen (D).

## Supplementary Index-2

### M Code for Sensitivity Analysis

~~~
function data = runsensitivity(modelobj, cs, variants)
% RUNSENSITIVITY calculate the sensitivities for SimBiology model,
% modelobj using the variant object(s), variants. Variant object
% settings supercede model property-values.
%
% To calculate the sensitivities of X with respect to Y, X is an output
% factor and Y is an input factor.
% Restore the configset after the task has run.
cleanup = onCleanup(@() restore(cs));
% Configure the configuration set for sensitivity analysis.
set(cs.SolverOptions, ‘SensitivityAnalysis’, true);
% Configure Normalization.
set(cs.SensitivityAnalysisOptions, ‘Normalization’, ‘none’);
% Configure the inputs.
input1 = modelobj.Compartments(1).Species(22);
input2 = modelobj.Compartments(1).Species(5);
set(cs.SensitivityAnalysisOptions, ‘Inputs’, [input1, input2]);
% Configure the output factors.
output1 = modelobj.Compartments(1).Species(44);
output2 = modelobj.Compartments(1).Species(45);
output3 = modelobj.Compartments(1).Species(16);
output4 = modelobj.Compartments(1).Species(47);
output5 = modelobj.Compartments(1).Species(34);
output6 = modelobj.Compartments(1).Species(60);
output7 = modelobj.Compartments(1).Species(9);
output8 = modelobj.Compartments(1).Species(8);
output9 = modelobj.Compartments(1).Species(33);
output10 = modelobj.Compartments(1).Species(24);
output11 = modelobj.Compartments(1).Species(55);
output12 = modelobj.Compartments(1).Species(11);
output13 = modelobj.Compartments(1).Species(42);
output14 = modelobj.Compartments(1).Species(52);
output15 = modelobj.Compartments(1).Species(57);
output16 = modelobj.Compartments(1).Species(21);
output17 = modelobj.Compartments(1).Species(51);
output18 = modelobj.Compartments(1).Species(30);
output19 = modelobj.Compartments(1).Species(54);
output20 = modelobj.Compartments(1).Species(36);
set(cs.SensitivityAnalysisOptions, ‘Outputs’, [output1, output2, output3, output4, output5, output6, output7, output8, output9, output10, output11, output12, output13, output14, output15, output16, output17, output18, output19, output20]);
% Run simulation.
data = sbiosimulate(modelobj, cs, variants, []);
% Plot the results.
plottype_Sensitivity_Matrix_Subplot(data, ‘<all>‘, ‘<all>‘);
plottype_Time(data, ‘<all>‘, ‘one axes’, ‘‘);
% ---------------------------------------------------------
function restore(cs)
% Turn off sensitivity analysis.
try
 set(cs.SolverOptions, ‘SensitivityAnalysis’, false);
 set(cs.SensitivityAnalysisOptions, ‘Inputs’, []);
 set(cs.SensitivityAnalysisOptions, ‘Outputs’, []);
catch ex
end
% ----------------------------------------------------------
function plottype_Sensitivity_Matrix_Subplot(tobj, input, output)
%SENSITIVITY_MATRIX_SUBPLOT Plots sensitivity Matrix using time integral.
%
% Plot the sensitivity matrix using the time integral for the sensitivities
% for the specified input and output factors.
%
% See also SBIOGETSENSMATRIX.
% Create a subplot if there is more than one run.
if (length(tobj) > 1)
 sbiosubplot(tobj, @sensubplotdata, output, input, false);
else
 sensubplotdata(tobj, output, input);
end
% ---------------------------------------------------------
function sensubplotdata(tobj, x, y)
% Parse the input arguments.
if strcmpi(y, ‘<all>‘)
 y = {};
end
if strcmpi(x, ‘<all>‘)
 x = {};
end
[t, R, states, inputs] = getsensmatrix(tobj, x, y);
[∼, in, out] = size(R);
result = zeros(in, out);
for i = 1:in
 for j = 1:out
  index = ∼isnan(R(:,i,j)) & ∼isinf(R(:,i,j));
  result(i,j) = trapz(t(index), abs(R(index,i,j)));
 end
end
% Plot the data.
if in == 1 || out == 1
 if out == 1 && in > 1
  hbar = barh(result);
  haxes = get(hbar(1), ‘Parent’);
  set(haxes, ‘ytick’, 1:length(states));
  set(haxes, ‘yticklabel’, states, ‘TickLabelInterpreter’,’none’);
  xlabel(sprintf(‘Sensitivity relative to: %s’, inputs{1}));
  ylabel(‘States’);
elseif in == 1
  hbar = bar(result);
  haxes = get(hbar(1), ‘Parent’);
  set(haxes, ‘xtick’, 1:length(inputs));
  set(haxes, ‘xticklabel’, inputs);
  ylabel(sprintf(‘Sensitivity of: %s’, states{1}));
  xlabel(‘Parameters’);
end
f = get(haxes, ‘Parent’);
title(‘Sensitivity - Time Integral’);
else
 hbar = bar3(result);
 view(2);
 title(‘Sensitivity Matrix - Time Integral’);
 zlabel(‘Sensitivity’);
 xlabel(‘Parameters’);
 ylabel(‘States’);
 haxes = get(hbar(1), ‘Parent’);
 set(haxes, ‘PlotBoxAspectRatioMode’, ‘auto’)
 f = get(haxes, ‘Parent’);
 while ∼strcmp(get(f, ‘type’), ‘figure’)
  f = get(f, ‘Parent’);
 end
 set(f, ‘Renderer’, ‘zbuffer’)
% Set the tick labels to the input and output names.
set(haxes, ‘xtick’, 1:length(inputs));
set(haxes, ‘xticklabel’, inputs);
set(haxes, ‘ytick’, 1:length(states));
set(haxes, ‘yticklabel’, states, ‘TickLabelInterpreter’, ‘none’);
% Color the bars by height.
for j = 1:length(hbar)
 zd = get(hbar(j), ‘zdata’);
 for i=1:6:size(zd,1)
  zd(i + [0 3 4], 2:3) = zd(i+1,2);
  zd(i + [1 2], [1 4]) = zd(i+1,2);
 end
 set(hbar(j), ‘cdata’, zd);
end
 colorbar;
end
% turn off default axes interactions and axes toolbar
set(f, ‘DefaultAxesToolbarVisible’, ‘off’);
allAxes = findobj(f, ‘-class’, ‘matlab.graphics.axis.Axes’);
set(allAxes, ‘Toolbar’, []);
arrayfun(@(ax)matlab.graphics.interaction.disableDefaultAxesInteractions(ax), allAxes);
% ----------------------------------------------------------
function plottype_Time(tobj, y, plotStyle, axesStyle)
%TIME Plots states versus time.
%
% TIME(TOBJ, Y, PLOTSTYLE, PROPS) plots the results of the simulation
% for the species with the specified Y versus time.
%
% If PLOTSTYLE is ‘one axes’ then data from each run is plotted into one
% axes. If PLOTSTYLE is ‘trellis’ then data from each run is plotted
% into its own subplot.
%
% If Y is ‘<all>’ then all data will be plotted.
%
% AXESSTYLE is a structure that contains axes property value pairs.
%
% See also GETDATA, SELECTBYNAME.
if ∼isempty(tobj(1).RunInfo.ConfigSet) &&
tobj(1).RunInfo.ConfigSet.CompileOptions.UnitConversion &&
all(strcmp({tobj.TimeUnits},tobj(1).TimeUnits)) && ∼isempty(tobj(1).TimeUnits)
 labelX = [‘Time (’ tobj(1).TimeUnits ‘)’];
else
 labelX = ‘Time’;
end
% Get the labels for the plot.
labelArgs = timeGetLabels(axesStyle, ‘States versus Time’, labelX, ‘States’);
if (length(tobj) > 1)
 switch (plotStyle)
 case ‘one axes’
  haxes = sbioplot(tobj, @timeplotdata, [], y, labelArgs{:});
 case ‘trellis’
  htrellis = sbiotrellis(tobj,@timesubplotdata, [], y, labelArgs{:});
  haxes = htrellis.plots;
end
% Configure the axes properties.
if isfield(axesStyle, ‘Properties’)
  set(haxes, axesStyle.Properties);
 end
else
 % Plot Data.
 handles = timesubplotdata(tobj, [], y);
 haxes = get(handles(1), ‘Parent’);
% Configure the axes properties.
if length(handles)>=1
 haxes = get(handles(1), ‘Parent’);
 if isfield(axesStyle, ‘Properties’)
  set(haxes, axesStyle.Properties);
 end
else
 haxes = [];
end
% Label the plot.
title(labelArgs{2});
xlabel(labelArgs{4});
ylabel(labelArgs{6});
% If Y is ‘<all>’ get all the data names.
if strcmpi(y, ‘<all>‘)
 [∼, ∼, names] = getdata(tobj);
else
 names = y;
end
% Create legend.
leg = legend(names, ‘Location’, ‘NorthEastOutside’);
set(leg, ‘Interpreter’, ‘none’);
% turn off default axes interactions and axes toolbar
if ∼isempty(haxes)
  f = get(haxes, ‘Parent’);
  set(f, ‘DefaultAxesToolbarVisible’, ‘off’);
  allAxes = findobj(f, ‘-class’, ‘matlab.graphics.axis.Axes’);
  set(allAxes, ‘Toolbar’, []);
  arrayfun(@(ax)matlab.graphics.interaction.disableDefaultAxesInteractions(ax), allAxes);
 end
end
%--------------------------------------------------------
function [handles, names] = timeplotdata(tobj, ∼, y)
colors = get(gca, ‘ColorOrder’);
numColors = length(colors);
% Preallocate handles if strcmpi(y, ‘<all>‘)
 [∼, ∼, names] = getdata(tobj(1));
else
 [∼, ∼, names] = selectbyname(tobj(1), y);
end
handles = zeros(length(names), length(tobj));
for i=1:length(tobj)
 % Get the data from the next run.
 nexttobj = tobj(i);
 % Get the data associated with y.
 if strcmpi(y, ‘<all>‘)
 [time, data, names] = getdata(nexttobj);
else
 [time, data, names] = selectbyname(nexttobj, y);
end
% Error checking.
if size(data,2) == 0
 error(‘Data specified do not exist.’);
end
set(gca, ‘ColorOrderIndex’, 1);
% Plot data. If there is only one state use different colors for runs.
if(size(data,2) ==1)
 hLine = plot(time, data, ‘color’,colors(mod(i-1,numColors)+1,:));
else
 hLine = plot(time, data);
end
 handles(:,i) = hLine;
end
% ---------------------------------------------------------
function handles = timesubplotdata(tobj, ∼, y)
% Get Data to be plotted.
if strcmpi(y, ‘<all>‘)
 [time, data] = getdata(tobj);
else
 [time, data] = selectbyname(tobj, y);
end
% Error checking.
if size(data,2) == 0
 error(‘Species specified do not exist.’);
end
% Plot Data.
handles = plot(time, data);
% ---------------------------------------------------------
function out = timeGetLabels(axesStyle, labelTitle, labelX, labelY)
out = {‘title’, labelTitle, ‘xlabel’, labelX, ‘ylabel’, labelY};
if isfield(axesStyle, ‘Labels’)
 allLabels = axesStyle.Labels;
if isfield(allLabels, ‘Title’)
 out{2} = allLabels.Title;
end
if isfield(allLabels, ‘XLabel’)
 out{4} = allLabels.XLabel;
end
if isfield(allLabels, ‘YLabel’)
  out{6} = allLabels.YLabel;
 end
end
~~~

